# Evolutionary trajectories are contingent on mitonuclear interactions

**DOI:** 10.1101/2022.09.11.507487

**Authors:** Damien Biot-Pelletier, Stefano Bettinazzi, Isabelle Gagnon-Arsenault, Alexandre K. Dubé, Camille Bédard, Tuc H. M. Nguyen, Heather L. Fiumera, Sophie Breton, Christian R. Landry

## Abstract

Critical mitochondrial functions, including cellular respiration, rely on frequently interacting components expressed from both the mitochondrial and nuclear genomes. The fitness of eukaryotic organisms depends on a tight collaboration between both genomes. In the face of an elevated rate of evolution in the mitochondrial genome, current models predict that maintenance of mitonuclear compatibility relies on compensatory evolution of the nuclear genome. Mitonuclear interactions would therefore exert an influence on evolutionary trajectories. One prediction from this model is that the same nuclear genomes but evolving with different mitochondrial haplotypes would follow distinct molecular paths towards higher fitness peaks. To test this prediction, we submitted 1344 populations derived from seven mitonuclear genotypes of *Saccharomyces cerevisiae* to more than 300 generations of experimental evolution in conditions that either select for a mitochondrial function, or that do not strictly require respiration for survival. Performing high-throughput phenotyping and whole-genome sequencing on independently evolved individuals isolated from endpoint populations, we identified numerous examples of gene-level evolutionary convergence among populations with the same mitonuclear background. Phenotypic and genotypic data on strains derived from this evolution experiment identify the nuclear genome and the environment as the main determinants of evolutionary divergence, but also show a modulating role for the mitochondrial genome exerted both directly and via interactions with the two other components. We finally recapitulated a subset of prominent loss-of-function alleles in the ancestral backgrounds and confirmed a generalized pattern of mitonuclear-specific and highly epistatic fitness effects. Together, these results demonstrate how mitonuclear interactions can dictate evolutionary divergence of populations with identical starting nuclear genotypes.

## Introduction

According to the endosymbiotic hypothesis, mitochondria derive from an ancestral bacterial symbiont that gradually specialized in its current cellular functions (Dyall et al. 2004; E.S. Fritsch et al. 2014). While most mitochondrial proteins are encoded in the nucleus, the vestigial genome of the organelle retains key genes thought to favor responsiveness to changing conditions (Allen et al. 2003; Lane and Martin 2010; Lane 2011). Gene products encoded in the two genomes are thus required for mitochondrial function. The mitochondrion is now the site of cellular respiration, and as such, plays a central role in energy production (Malina et al. 2018). Other important metabolic processes involve the mitochondrion, including the biosynthesis of heme (Ferreira 1995), iron-sulfur clusters (Read et al. 2021) and steroids (Jordá and Puig 2020). Mitochondria are also involved in nutrient sensing, the regulation of redox homeostasis, and several signaling pathways (Malina et al. 2018).

Components encoded in the nuclear and mitochondrial genomes interact extensively, sometimes encountered together as part of mitochondrial protein complexes (Malina et al. 2018). Cytochrome *c* oxidase (Cooper et al. 1991) and the cytochrome *bc1* complex (Hunte et al. 2003), components of the respiratory electron transport chain, are assembled from proteins encoded by both genomes. In yeast, the mitogenome encodes for mitochondrial ribosomal RNAs and one ribosomal polypeptide, while most protein components are found in the nuclear genome (Desai et al. 2017). In humans, the nucleus-encoded protein LRPPRC is required for cytochrome *c* maturation via interaction with mitochondrial mRNAs, and mutations that alter this function are associated with severe disease (Xu et al. 2012). These examples illustrate the critical importance of mitonuclear interactions, and how their disruption may severely reduce fitness.

Yet, because they are distinct and segregated to separate cell compartments, molecular evolution in the nuclear and mitochondrial genomes is partially uncoupled. Mutation rate tends to be higher in mtDNA (Fritsch et al., 2014; Lynch et al., 2008), especially in organisms with uniparental inheritance and lacking a dedicated mitochondrial mismatch repair system (Mason and Lightowlers 2003; Havird and Sloan 2016). In many organisms, uniparental inheritance also leads to smaller effective population size and the virtual lack of recombination (Lynch et al. 2006; Hagström et al. 2014). More generally, mitochondria evolve in a regulated microenvironment – the cytoplasm with its nuclear supply of mitochondrial proteins – that differs from the extracellular milieu affecting the entire cell. Illustrating this peculiar aspect of mitochondrial evolution is the suppressivity of ρ^-^ mitogenomes in yeast. In these cells, despite a strongly deleterious effect on organismal fitness, copies of the mitochondrial genome deleted at loci essential for respiration may outcompete healthy copies thanks to their higher replication origin density (de Zamaroczy et al. 1981; Mangin et al. 1983; Bernardi 2005). Furthermore, deletion screens in yeasts have also repeatedly identified signals of frequent essentiality switching among genes involved in mitochondrial functions, both across (Kim et al. 2010) and within species (Caudal et al. 2022; Chen et al. 2022), suggesting enhanced evolvability at these loci. In summary, because of unique characteristics and selection pressures, the rate of evolution tends to be higher in the mitochondrial genome than in the nuclear genome, with a critical influence on genetic divergence, speciation, reproductive strategies, and morphology (Hill 2015).

A model has emerged in which rapid and essentially non-adaptive changes to the mitochondrial genome exert a constant selective pressure on the nuclear genome to accumulate compensatory mutations (Dowling 2014). Rapid changes in mtDNA would drive the nucleus towards increasing divergence, the evolution of Dobzhansky-Muller incompatibilities favoring speciation via post-zygotic reproductive isolation (Freel et al. 2015). Such incompatibilities are expected to arise following hybridization between allopatric populations, whose mitonuclear genotypes have evolved independently under different environmental conditions. As extensively described in the copepod *Tigriopus californicus*, hybrids may display fitness loss and reproductive incompatibility when initially separated populations come into contact, generating mismatched mitonuclear genotypes (Burton and Barreto 2012; Barreto et al. 2018; Burton 2022). A growing body of evidence accumulates in favor of this model. Early studies showed that species divergence is associated with a loss of mitonuclear compatibility (Niki et al. 1989; Osuský et al. 1997; Dey et al. 2000a; McKenzie and Trounce 2000; Špírek et al. 2000; Yamaoka et al. 2000; Kenyon and Moraes 2002). More recent reports provide examples of reproductive isolation and hybrid fragility associated with mitonuclear incompatibilities (Lee et al. 2008; Chou et al. 2010; Paliwal et al. 2014; Ma et al. 2016; Haddad et al. 2018). The power of mitochondrial markers and genome structures as phylogenetic predictors further supports a driving role for the evolution of mtDNA in divergence and speciation (Robba et al. 2006; Hendrich et al. 2010; Wolters et al. 2015).

Current understanding about the cell and molecular biology of the mitochondrion owes much to methods and approaches pioneered in the yeast *S. cerevisiae* (Rutter and Hughes 2015). As a facultative anaerobe, this yeast can be explicitly selected for respiratory function, via the use of fermentable and non-fermentable carbon sources. This facilitates cytoduction, in which mitochondrial genomes are exchanged between individuals of the same or different species, generating mitonuclear hybrids (Zakharov and Yarovoy 1977; Barrientos et al. 1998). Cytoduction and related methods have been applied to the study of evolutionary questions in mitochondrial biology, for example investigating the compatibility of mitochondrial genomes between closely related species. Coherent with a role for mitochondria in defining the interspecific barrier, these studies showed a rapid decline in the efficiency of oxidative phosphorylation as species divergence increases between the nuclear and mitochondrial parents of mitonuclear hybrids (Barrientos et al. 1998; Dey et al. 2000a; Dey et al. 2000b; McKenzie and Trounce 2000; Yamaoka et al. 2000; Kenyon and Moraes 2002; Lee et al. 2008; Ma et al. 2016). Cytoduction has also helped identify key genes and molecular functions involved in the mitonuclear incompatibilities that arise between yeast species (Spirek et al. 2015). Collections of yeast mitonuclear hybrids have recently helped explore the genetic space of mitonuclear interactions, revealing the impact of mitonuclear interactions on fitness, and the complex third-order relationships that exist between environmental conditions, the nuclear genome, and the mitochondrial genome (Paliwal et al. 2014; Vijayraghavan et al. 2019; Nguyen et al. 2020).

Standing models thus predict that mitonuclear interactions must influence the path of evolution. Available evidence from wild individuals and populations supports this model, yet direct experimental testing of this hypothesis in naturally evolving populations is challenging. Experimental evolution in microbes is ideally suited to address this problem, because it can be replicated, enabling the observation of repeatable patterns among identically but independently evolved individuals and populations. Indeed, one of the fundamental questions that underlie much of the experimental evolution literature concerns the predictability of evolution. Given identical genetic and environmental circumstances, will independently evolving populations take the same evolutionary path? Will phenotypes be identical, will the same alleles be selected, or at least will the same genes and functional systems be affected? Repeatable outcomes have been reported in terms of both phenotype and genotype (Wichman et al. 1999; Cooper et al. 2003; Mcdonald et al. 2009; Herron and Doebeli 2013; Lang et al. 2013). Available evidence does not decisively argue for predictability (Bull and Molineux 2008; Łuksza and Lässig 2014), yet the controlled context of experimental evolution seems to lead to evolutionary convergence at the functional complex and gene levels (Tenaillon et al. 2012). Another important advantage of experimental evolution in microbes is the conservation of evolving populations at different steps of the evolutionary process. Evolved individuals can thus be compared experimentally to their ancestors. Mutations selected by evolution can be reconstituted in ancestral backgrounds and assessed for their impact on fitness using marker-assisted competition assays (Breslow et al. 2008; Lang et al. 2013), allowing the precise dissection of the genetic architecture of evolved mutants. Building on these methodological foundations, experimental evolution studies have demonstrated that the path to adaptation is contingent on the genetic makeup of the evolving population and the characteristics of its environment (Blount et al. 2008; Meyer et al. 2012). Therefore, analysis of evolutionary convergence in replicated experimental evolution setups appears as a mean to assess the impact of specific environmental and genetic factors, such as mitonuclear interactions, on evolutionary trajectories and outcomes.

In this study, we leverage a collection of mitonuclear hybrids from *S. cerevisiae* (Paliwal et al. 2014; Wolters et al. 2018; Nguyen et al. 2020) previously reported to display mitonuclear interactions in a wide array of conditions (Nguyen et al. 2020). To investigate the evolutionary impact of these interactions, we identified example strains from the collection that showed reduced fitness upon mitochondrial replacement. We performed experimental evolution on 1344 replicate populations of the identified strains, varying the intensity of selection on mitochondrial function in an experiment spanning more than three hundred generations. Evolving mitonuclear hybrids provided accelerated mimics of the evolutionary dynamics encountered in organisms with rapidly diverging mitogenomes. The level of replication provided us with a wealth of identically yet independently evolved individuals. We predicted that high-throughput growth assays and whole-genome sequencing on these individuals would allow us to detect patterns of evolution influenced by mitonuclear interactions, at the phenotypic or genotypic levels.

## Results

### Mitonuclear interactions in yeast cybrids

Our founding intent was to study the influence of mitonuclear interactions on the path of evolution. We therefore screened a collection of mitonuclear hybrids of *S. cerevisiae* for evidence of such mitonuclear interactions. This collection contains 225 cybrid strains of yeast generated by cytoduction in four replicates and representing all possible combinations of nuclear and mitochondrial genomes from 15 trains of *S. cerevisiae* (Nguyen et al. 2020). Though widespread mitonuclear interactions were previously found within the collection, we screened it for additional evidence, recording growth rate for all strains as a proxy for fitness in a specific condition – liquid medium with glycerol as the sole carbon source – that requires mitochondrial activity to sustain facultative anaerobic yeast. As observed previously (Nguyen et al., 2020; Paliwal et al., 2014; Wolters et al., 2018), the screen showed the predominant effect of the nuclear genome on fitness but revealed a modulating influence from the mitochondrial genome (**Fig S1**). Some pairs of nuclear and mitochondrial parents displayed a depression in fitness upon cytoduction, potentially indicating mismatched nuclear and mitochondrial genomes. Prominently, mitonuclear hybrids formed between strain DBVPG6044 (D) and strains 273614N (N) and Y12 (Y) were submitted to further scrutiny to formally test the phenotypic influence of the nuclear and mitonuclear genomes and their interactions (**Fig 1, S2**). Following a two-letter nomenclature, indicating the nuclear background first and the mitochondrial background second, genotypes of interest were: NN, ND, DN, DD, DY, YD, and YY (**Fig 1A**). These genotypes can be regarded as two sets of mitonuclear swaps: the N<->D set consisting of genotypes NN, ND, DN, DD; and a D<->Y swap consisting of DD, DY, YD, YY. We estimated fitness of the strains by recording their growth rate and carrying capacity in two environments: fermentable (FF) and non-fermentable (NF) media, containing glucose and glycerol as the sole carbon source, respectively. For the two sets of mitonuclear swaps, we drew interaction plots and performed two-way ANOVAs to test the influence of both genomes and their interactions on fitness (**Fig 1B**, **S2**). Consistent with previous observations, the nuclear genome appeared as the main determinant of fitness, significantly affecting growth in all conditions and for all growth parameters, with the mitochondrial background playing but an occasional role. Notably, we noted that growth was dependent on statistically significant mitonuclear interactions in both sets. We also assayed the activity of a panel of enzymes associated with the mitochondrion in whole cell lysates of the same strains grown in non-fermentable medium (**Fig S3**). Independent effects of nuclear and mitochondrial genomes and interacting mitonuclear effects varied depending on the enzyme assays and strain set. Significant effects for mitonuclear interactions were detected for malate dehydrogenase (N<->D) (**Fig 1C**), citrate synthase (N<->D), and ATPase (D<->Y) (**Fig S3**). We thus accumulated a body of evidence demonstrating mitonuclear interactions by showing that interrupting coevolved mitonuclear genotypes alters fitness in strains N, D, Y and their cybrid derivatives. This enabled further studies on the evolutionary influence of mitonuclear interactions in yeast.

**Figure 1.**
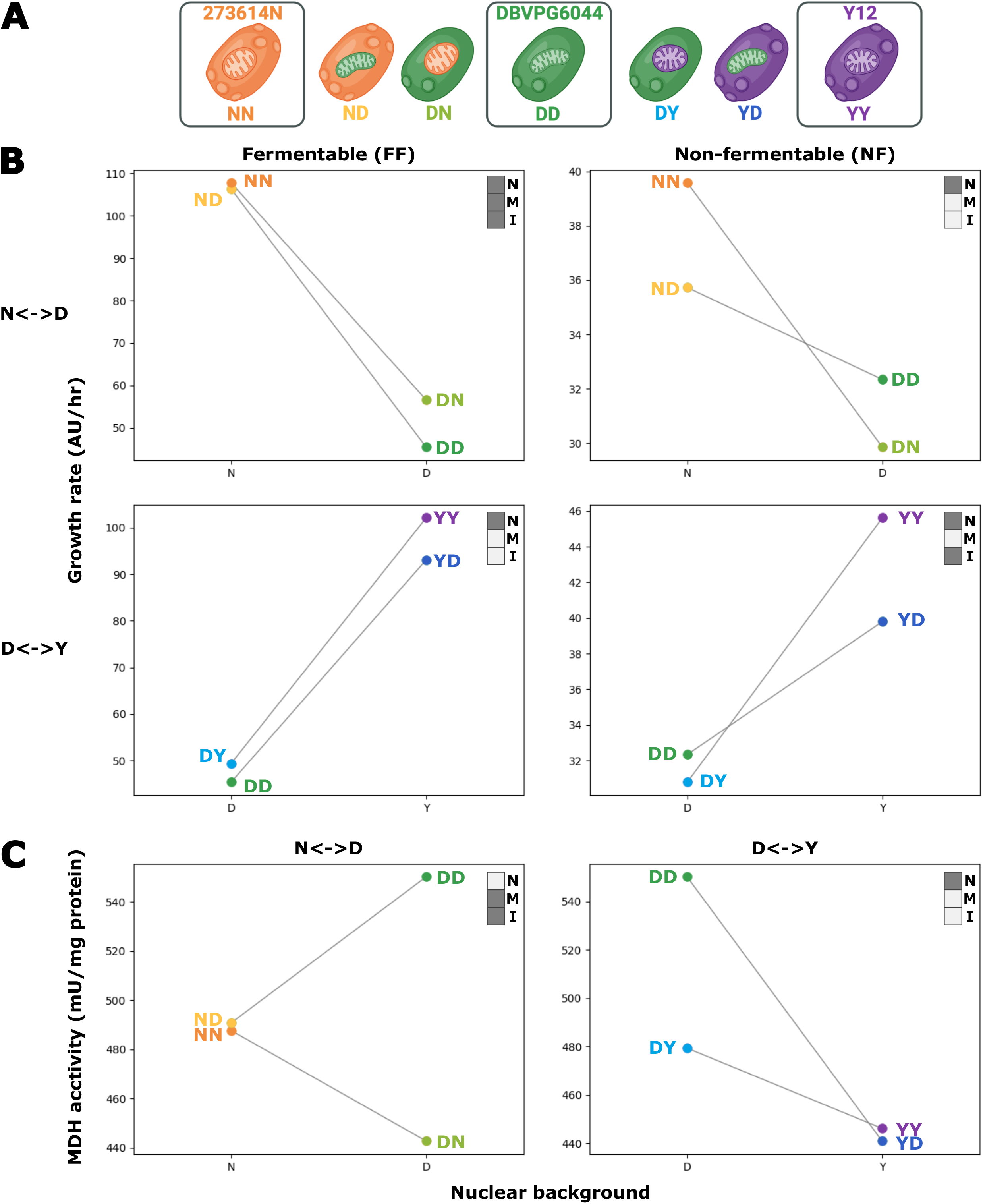
Adaptive evolution of mitonuclear hybrids. **(A)** Yeast strains submitted to experimental evolution were derived from strains 273614N (NN), DBVPG6044 (DD) and Y12 (YY). Cytoduction of NN and YY with DD yielded two cybrids each. (ND and DN, DY and YD). We thus evolved seven ancestral mitonuclear backgrounds. **(B)** Each of these backgrounds was evolved in fermentable (FF) and non-fermentable (NF) media, as 96 replicates. Evolving populations were propagated for approximately 300 generations by daily robot-assisted serial passage. At the end of the experiment, a single independently evolved individual was isolated from each population. Evolved individuals were assessed for growth rate in fermentable **(C)** and non-fermentable **(D)** media, showing significant gains in fitness in most cases. Asterisks indicate median fitness in evolved strains significantly higher than ancestors (Mann-Whitney U-test p-value < 0.05).

### Yeast mitonuclear hybrids submitted to experimental evolution undergo fitness changes influenced by mitonuclear interactions

We evolved 96 populations for each of the seven mitonuclear backgrounds derived from strains N, D and Y. This experiment was performed in conditions that require or do not require mitochondrial respiration using non-fermentable and fermentable sugars, respectively. We cultured 1344 independently evolving populations by daily serial passage for 58 days, amounting to approximately 300 generations (**Fig 2A**). Frozen stocks were prepared at the beginning and end of the experiment, as well as every seventh passage. A subset of 230 populations went extinct or were terminated because of bacterial contamination, while 1114 populations survived to the end. A single individual was isolated by streaking for single colonies from stocks of each of the surviving populations.

**Figure 2.**
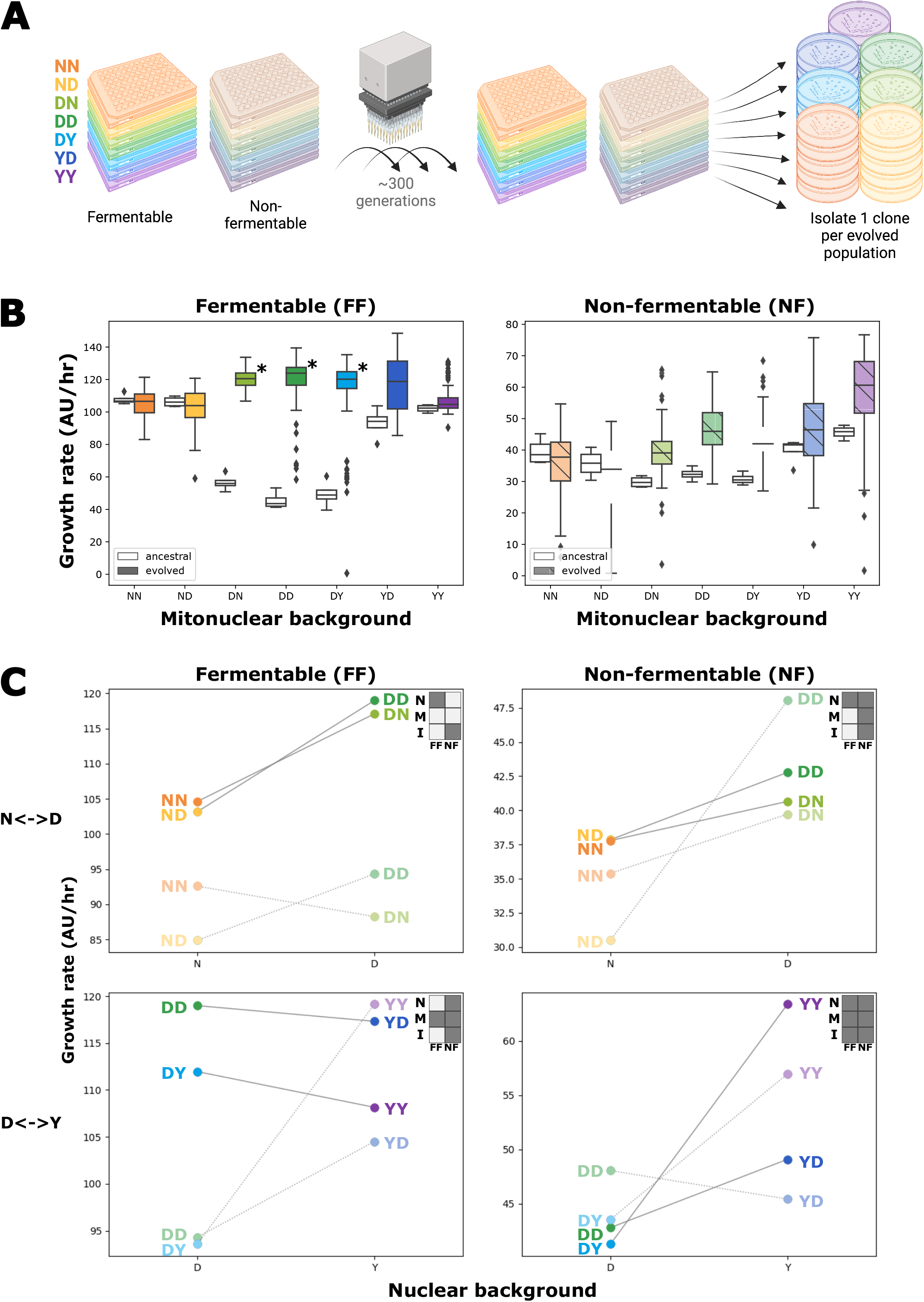
Whole-genome sequencing reveals mitonuclear background- and carbon source-specific genomic changes in experimental evolution isolates. **(A)** Fraction of sequenced bases mapped to the mitochondrial genome was used as a proxy for the proportion of DNA belonging to the mitochondrial genome, revealing a mitonuclear background-specific impact for both cytoduction and evolution. **(B)** Changes in chromosome copy numbers (aneuploidies), and smaller, gene-scale copy number variants (CNVs) we detected using depth of coverage information. Many single-nucleotide variants (SNVs) were also called. All those changes were unevenly distributed among mitonuclear backgrounds and carbon sources.

We next measured the extent of adaptation in evolved individuals by recording growth on both fermentable and non-fermentable media, in an automated and highly replicated manner (**Fig S4**). As expected, the evolved isolates generally displayed median fitness above ancestral values, but the magnitude of the gains was dependent on the fitness of the founding strains (**Fig 2B**). Best illustrating this trend are strains carrying the D nuclear genome for which comparingly low initial fitness in fermentable media represented potential for large gains (**Fig 2B**). In comparison, mean fitness gains in fermentable medium are modest if not null in backgrounds carrying the Y and N nuclei. Interestingly, a similar endpoint fitness, close to the value of the fittest ancestors (NN, YY) is reached across backgrounds in fermentable medium suggesting that evolved clones are, on average, approaching fitness optimum in this environment. Evolution in non-fermentable medium also generally led to higher median fitness. Here also, the N nuclear genome is associated with the lowest median fitness among evolved individuals. For example, in fermentable medium, NN and ND individuals evolved in non-fermentable conditions generally appear less fit than their ancestors. This further indicates that, in fermentable medium, these strains are close to fitness optimum, where most mutations are expected to be deleterious, including those that are adaptive in other environments. Of note, in non-fermentable medium, the fittest of the ancestral backgrounds (YY) also displays one of the largest median gains in fitness, suggesting individuals evolved in this environment are still largely far from fitness optimum.

All evolved individuals were assayed for growth in fermentable and non-fermentable media. Gains achieved through evolution in one carbon source often translated into gains in the other. Indeed, in most mitonuclear backgrounds, individuals evolved in fermentable and non-fermentable medium display similar fitness in both carbon sources (**Fig S4**). Strikingly, in non-fermentable conditions, strains evolved in this medium are frequently matched or outperformed by those evolved in fermentable medium (**Fig S4BD**), as they likely spent a portion of each evolution cycle respiring in the stationary phase. Similarly, in fermentable medium, while non-fermentable-evolved individuals tend be less fit than their fermentable-evolved counterparts, they still demonstrate significant fitness gains over ancestral levels (**Fig S4AC**). These observations suggest that much of the gain in fitness is related to aspects common to both environments, either in terms of media composition or culture conditions.

We used the fitness of evolved individuals to test the effect of mitonuclear interactions on evolutionary outcomes. We drew interaction plots and performed two-way ANOVAs, revealing patterns of relative fitness that bore both resemblances and differences with those of ancestral strains (**Fig 2C**, **S5**). Nuclear effects on fitness are most frequent in both ancestral and evolved strains. However, evidence for the influence of the mitochondrial background and mitonuclear interactions is more frequent in fitness data of the evolved strains. These results show an influence for mitonuclear interactions on the outcomes of experimental evolution at the phenotype level.

### Sequencing of evolved genomes reveals patterns of evolutionary convergence

To probe the molecular mechanisms involved in the adaptation of evolved isolates, we sequenced each of their genomes, along with that of their ancestors, aligning to published sequences of ancestral nuclear and mitochondrial backgrounds. Our partially redundant dataset comprised 1505 non-empty sequencing libraries and provided genomic data for 1114 evolved isolates and 28 ancestral strains. With a 31x median depth of coverage, most libraries were suitable for the confident calling of mutations, from the whole-chromosome to single-nucleotide scales (**Data S1**). We first examined chromosome-scale changes, beginning with the mitochondrial genome. In contrast with published methods that rely on median depth at stably covered loci *COX1* or *COX3* (Puddu et al., 2019), and acknowledging the complex and highly branched topology of *S. cerevisiae* mtDNA (Bendich 1996; Bendich 2007), we abstained from claims about mitogenome copy number, and used the fraction of sequence bases mapped to coding regions of the mitochondrial genome to probe general changes to mtDNA, including structure, copy number and others, incurred via experimental evolution and cybridization (**Fig 3A**). Because of disparities in the sequence, structure and quality of the reference mitochondrial genomes, we restricted comparisons to strains that share the same mitochondrial background. We observed changes in mtDNA among the ancestral mitonuclear hybrids, echoing enzyme assays (**Fig S3**) hinting to changes in mitochondrial metabolism upon cytoduction. We also observed a trend towards a decrease in the fraction of coding mtDNA in evolved isolates. This change is likely not the result of widespread petite genotypes, since most isolates display fitness in non-fermentable media that is equal or superior to their non-evolved ancestors (**Fig S4BD**). Similarly, extent of the contraction in mtDNA fraction is dependent on both mitonuclear background and evolution conditions. Because environmental demands on respiratory metabolism affect mtDNA copy numbers, our observations about mtDNA fraction are chiefly relevant to the fermentable conditions in which we propagated the yeast used for gDNA extraction (Galeota-Sprung et al., 2022).

**Figure 3.**
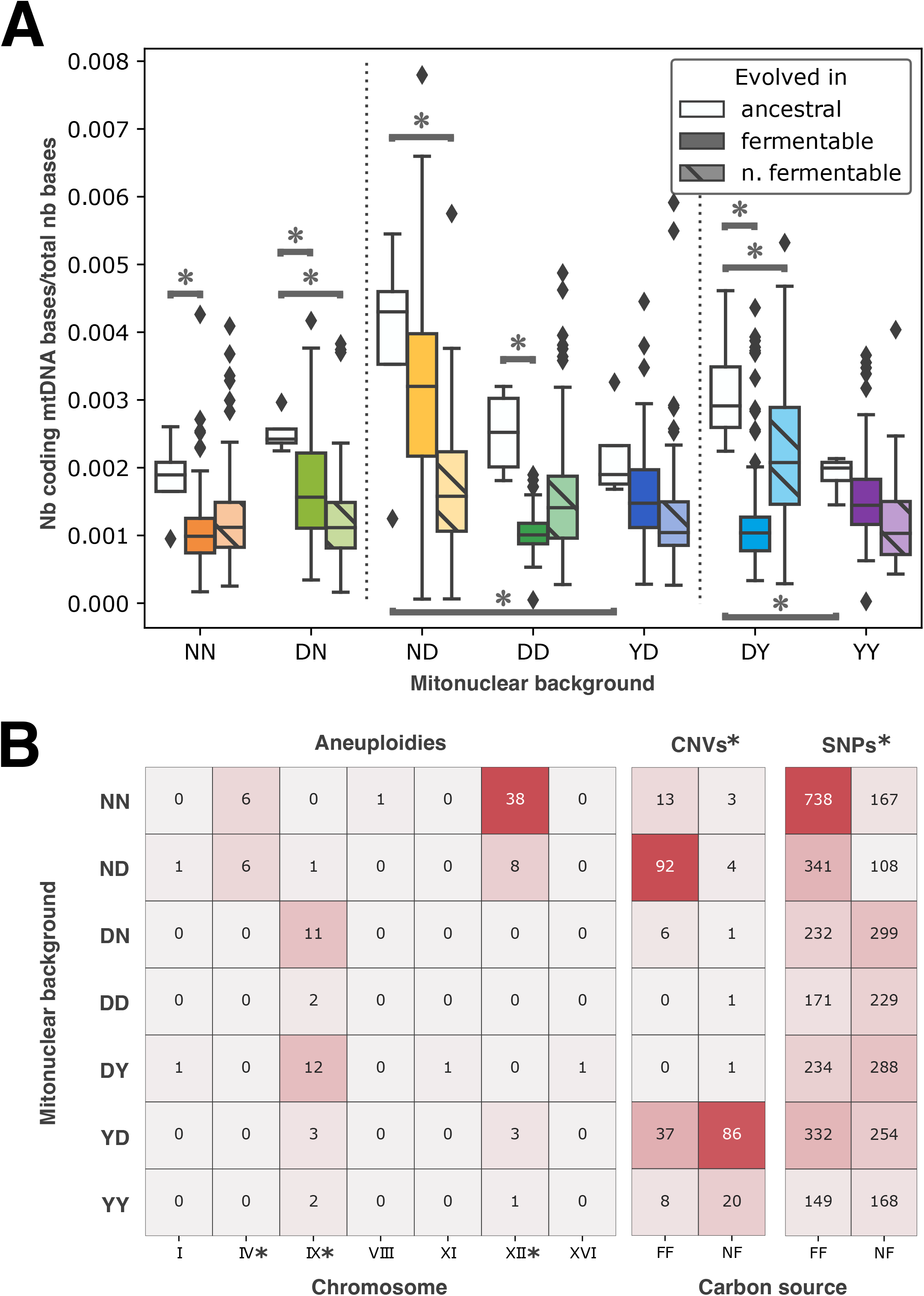
Mutation at certain loci is associated with quantitative phenotype changes. Mutant loci associated with quantitative changes in phenotype were identified by performing two statistical tests. Scatter plots show on the x-axis the negative log p-value that individual mutants at a given locus have phenotype identical to their parents, and on the y-axis, the negative log p-value that a phenotypic effect of this size could be observed from a set of random individuals of equal size. Dotted lines indicate significance thresholds for p-values > 0.05, corrected for false-discovery rate. Color of the data points indicates the mitonuclear background most often affected by mutation at the locus, while shape indicates carbon source in which the locus was most often mutant. This analysis was performed for carrying capacity **(A, B)** and growth rate **(C, D)**, in fermentable **(A, C)** and non-fermentable **(B, D)** media, as well as for the fraction of mtDNA per cell **(E)**.

We detected copy number increases for several chromosomes, prominently including chromosomes XII, IV and IX (**Fig 3B**, **Fig S6**). We detected aneuploidies at chromosome XII in 50 independently evolved strains: these changes were significantly enriched in presence of the N nuclear genome and the matching mitochondrial genome. We observed a similar pattern for aneuploidies of chromosome IV. We most often found extra copies of chromosome IX in mitonuclear hybrid derivatives of strain D (**Fig 3B**). Along with changes in mtDNA, these repeated large-scale genomic changes suggested a pattern of convergent evolution influenced by the identity and interaction of nuclear and mitochondrial genomes.

We next identified smaller scale genetic changes. Following stringent bioinformatics pipelines, we called 272 copy number variants (CNVs), ranging in size from 184 bp to 1091 bp, and 3710 single nucleotide variants (SNVs) (**Data S3**). Neither CNVs nor SNVs were distributed randomly among mitonuclear backgrounds and carbon sources (chi^2^ test p-values < 0.05). This fact may appear most obvious observing that only nine CNV calls were made in nuclear derivatives of strain D. In stark contrast, ND derivatives evolved in fermentable medium and YD derivatives evolved in non-fermentable medium accumulated 92 and 86 CNVs, respectively (**Fig 3B**). One may note that both backgrounds derive from mitochondrial swaps. In the same way, SNVs are unevenly distributed, yet the large number of calls ensures that more than a hundred mutations were detected per mitonuclear background in each carbon source. Together, these small-scale changes provided further hints that both mitonuclear interactions and the environment inflected adaptive trajectories in our mitonuclear hybrids.

### Dissimilar mutational profiles provide evidence for mitonuclear specific evolutionary trajectories

From the distribution of aneuploidies and CNVs we hypothesized that each mitonuclear background and each carbon source could be associated with at least partially distinct sets of mutations. We tested this hypothesis by calculating Bray-Curtis dissimilarity between the sets of mutations accumulated by each background in each environment (**Fig S7**). While variable, mean dissimilarity between the mutational profiles of all pairs of evolutionary conditions is high at 0.93. To better grasp the influence of carbon sources and nuclear and mitochondrial backgrounds on mutational profiles, we used the dissimilarity matrix as input for non-parametric multi-dimensional scaling (NMDS, **Fig 4A**). Strikingly, this analysis reveals a broad pattern in which backgrounds evolved in the same environment cluster together along a single axis. A shared nuclear background is also associated with smaller distances and a tendency to cluster together. This analysis also suggests that the mitochondrial background more modestly influences mutational divergence. Visual examination of the dissimilarity matrix (**Fig S7**) also suggested that a shared nuclear background or evolution in the same environment could be associated with reduced dissimilarity in mutational profile. We formally tested these intuitions by partitioning the dissimilarity matrix according to match or mismatch with respect to environment, nuclear background, and mitochondrial background (**Fig 4B**). We performed a three-way ANOVA on the data partitioned in this way, revealing a significant effect on dissimilarity for all three matches and their interactions. Lowest dissimilarities are associated with pairs that display a single mismatch, either at the mitochondrial or environmental levels, whereas mismatched nuclei suffice to maximize dissimilarity. Highest dissimilarity is observed when both the environment and the nucleus differ, regardless of the mitochondrion. Together, these results confirm NMDS outcomes by ascribing a major role in mutational divergence to the nuclear background and identifying the environment as more impactful than the mitochondrial background. Critically, all three aspects impact evolutionary divergence and interact to yield dissimilar mutational profiles.

**Figure 4.**
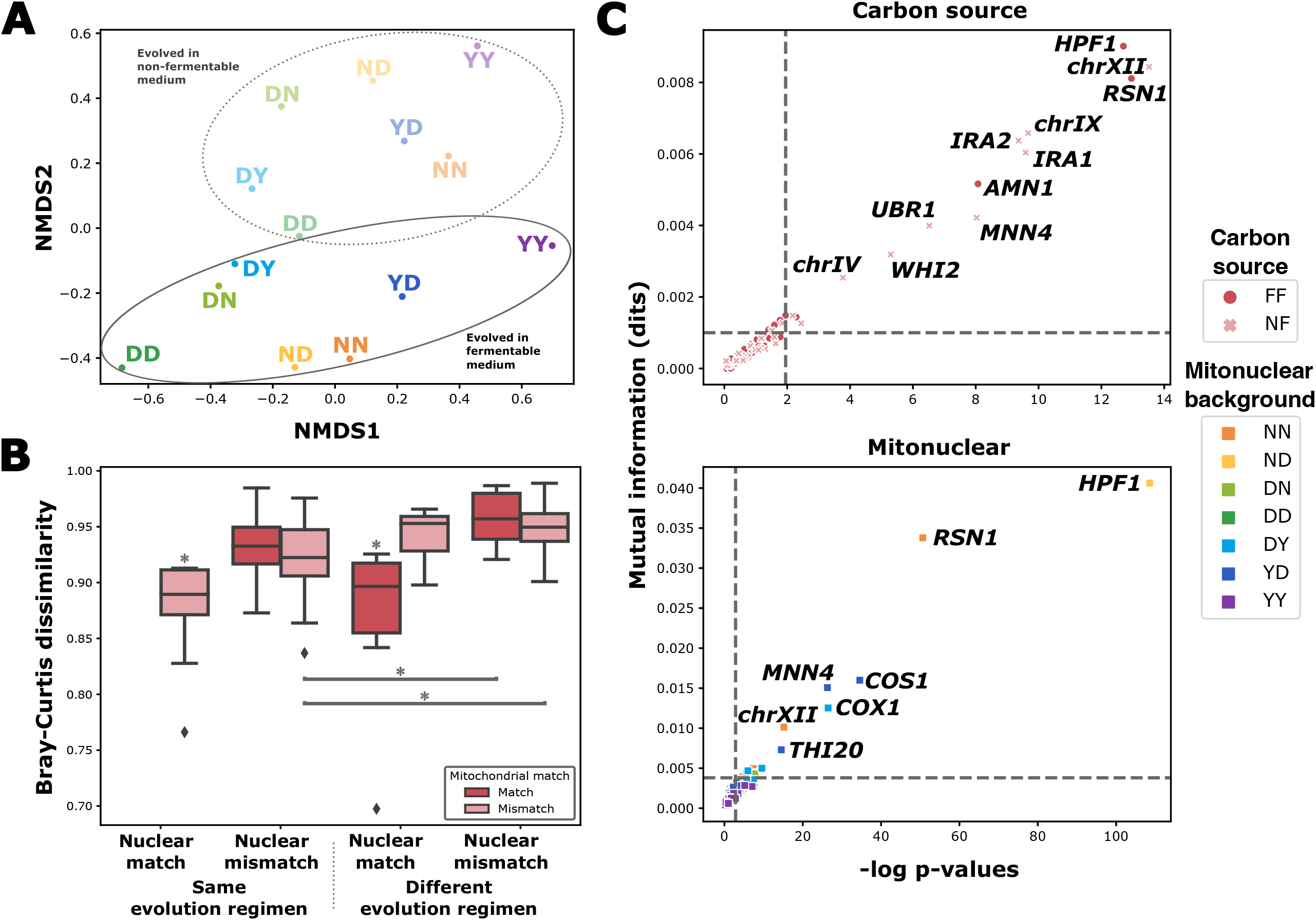
Mutations display convergence that is carbon source and mitonuclear specific. Loci were considered specific for a certain carbon source **(A)** or mitonuclear background **(B)** if their distribution among categories differed significantly from that of all detected mutations, taken as a whole. Chi^2^ tests were performed on contingency tables of all mutations, classifying them as mapping or not to the locus, and according to either carbon source or mitonuclear background. Mutual information was calculated from the same contingency tables. Specificity thresholds, indicated by dotted lines, were set at a p-value of 0.05, corrected for false-discovery rate, and at two standard deviations above mean (∼95^th^ percentile) mutual information across all loci.

We next asked if patterns of specificity were observed at the gene level and examined annotations associated with CNV and SNV calls (**Data S4**). Like aneuploidies, gene-level annotation indicated evolutionary convergence at several loci. Indeed, hundreds of genes are associated with more than one mutation among evolved clones, and 43 display ten or more. Along with aneuploidy at chromosome XII, nuclear genes *RSN1, HPF1, MNN4, AMN1* and mitochondrial gene *COX1* are the annotations associated with the strongest signals of evolutionary convergence, with tens to more than a hundred calls per locus mapping to these areas of the genome. Examining mutation counts associated with these loci in each mitonuclear background and carbon source (e.g. **Fig S8**), we detected several examples of potential specificity.

Our survey of genetic changes in evolved individuals, at all scales examined, thus provided evidence for evolutionary convergence within a given mitonuclear background, and strongly suggested that adaptive changes followed a pattern of specificity to both mitonuclear background and environmental conditions. To test this hypothesis, for each annotation, we performed chi^2^-tests of independence on contingency tables partitioning all mutations according to their association with the annotation, and according to either mitonuclear background or carbon source. We also computed mutual information from the same contingency tables. Both a significant p-value from this test and significantly above average mutual information were required to identify specific annotations (**Fig 4C**). Similar computations were performed, partitioning mutation counts according to nuclear and mitochondrial backgrounds alone, or conjunctions of background with carbon source (**Fig S9**). These analyses revealed areas of the yeast genome biased towards de novo mutation in a mitonuclear background- and carbon source-specific manner. For example, out of 50 aneuploidies detected at chromosome XII, 38 were found in background NN, and they appear to have been selected almost exclusively in non-fermentable medium. Similarly, 94% of the 88 CNVs at gene *HPF1* were selected in background ND evolving in fermentable medium (**Fig S8**). From this data and our earlier analysis of mutational dissimilarity, we concluded that interactions of the environment with both nuclear and mitochondrial backgrounds inflected evolutionary trajectories at the molecular level.

### Reconstituting loss-of-function mutations in ancestral backgrounds leads to mitonuclear background-specific fitness effects

Collected evidence suggested each mitonuclear background accumulated a distinct set of mutations during adaptive evolution. Therefore, we hypothesized that the phenotypic effects of individual mutations selected during the experiment were epistatic with the mitonuclear background. We tested the statistical association between mutant annotations and available phenotype measurements and found several candidate causal loci (**Fig S10**). From this list of loci, patterns of convergent evolution and both mitonuclear and carbon source specificity, we identified a core set of annotations potentially involved in adaptation or otherwise implicated in the evolution of our mitonuclear hybrids. Mapping of mutations on these annotated features and prediction of their mutational effects revealed widespread stop codons alongside missense mutations at conserved positions (e.g., **Fig S11**). We therefore hypothesized that loss-of-function at these and other loci was a widespread adaptive mechanism selected by our experiment, as observed for other experimental populations evolved in constant environments (Kvitek and Sherlock 2013).

To mimic the effect of loss-of-function alleles, we generated a replicated set of knockouts in fluorescent derivatives of each of the seven ancestral backgrounds at eight loci identified in our dataset. Loci were chosen based on frequency of mutation, statistical association with changes in phenotype, technical feasibility, gene essentiality, and both mitonuclear and carbon source specificity. We competed the knockouts against their wild type counterparts in the conditions of the evolution experiment. From these competitions, we estimated the fitness effect of loss-of-function at these loci in all mitonuclear backgrounds and both fermentable and non-fermentable media (**Fig 5**). Loss-of-function at these loci led in most cases to significant gains in fitness, validating their roles in adaptive evolution. The magnitude of the changes in fitness associated with each knock-out differed significantly between mitonuclear backgrounds and was influenced by nuclear and mitochondrial backgrounds as well as mitonuclear interactions (two-way ANOVA p-value < 0.05, **Fig S12**). Some loci even showed opposite effects between different mitonuclear backgrounds, further underlining the strongly epistatic nature of their impact on fitness. This result provided additional support for a decisive role of mitonuclear interactions in shaping and inflecting the path of adaptive evolution.

**Figure 5.**
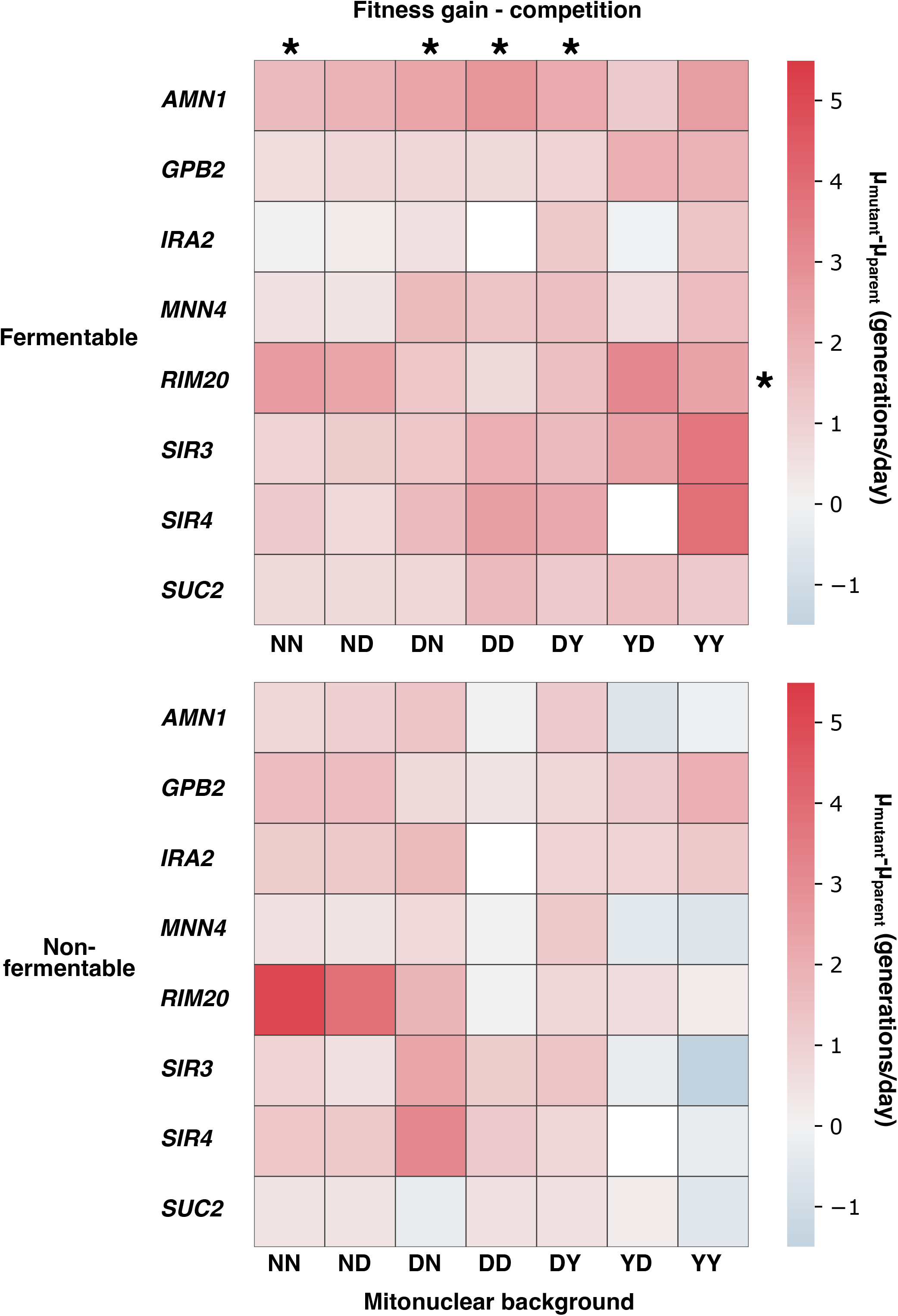
Fitness effect for loss-of-function at select loci is dependent on mitonuclear background and carbon source. Gene knockouts were performed at the indicated loci in fluorescent-derivatives of all seven ancestral strains used in the evolution experiment, mimicking loss-of-function alleles. These knockouts were competed against their deletion-free counterparts of the same mitonuclear genotype in fermentable (**top**) and non-fermentable (**bottom**) media. This enabled estimation of the fitness effect of loss-of-function at these loci, reported in shades of red (positive effect) or blue (negative effect). For each locus, effect of deletion was found to differ significantly between mitonuclear genotypes (ANOVA p-value < 0.01). Similarly, loci had differing effects on any given mitonuclear genotype (ANOVA p-value < 0.01). Fitness effect of each locus in across mitonuclear genotypes did not correlate with frequency of mutation at the locus in each genotype, with one exception indicated by an asterisk on the right-hand side (spearman rho > 0.95). Asterisks at the top of heatmaps indicate genotypes that display rank order correlation between locus fitness effect preference and frequency of mutation at the locus (spearman rho > 0.7, p-value<0.05 that slope of regression =0).

Competition assays provided additional support for evolution contingent on mitonuclear interactions. Yet, for most tested loci, differential fitness effect patterns did not match predictions derived from our analysis of evolutionary convergence (**Fig 5, Fig S13**). Contrary to expectation, apart from *RIM20* knockouts growing in fermentable medium, rank correlation was poor between estimated fitness effects and mutation frequency in evolved mutants (Spearman correlation p-value on the slope > 0.05). In other words, frequent mutation at a locus does not reliably predict strong gains in fitness in knockouts of that locus. In contrast, in fermentable medium, predictions of locus preference based on mutation frequency matched fitness effect-based predictions of preference, for half of our mitonuclear backgrounds (**Fig 5, Fig S13**).

We explored possible explanations for the partial disconnect between convergence- and fitness effect-based predictions. The experiment was based on the simplifying assumption that complete loss-of-function is the primary and sole mechanism underlying the fitness effect of mutation at all examined loci. Different or concurring mechanisms are likely involved and may explain some of the discrepancy between fitness data and our analysis of evolutionary convergence. Such mechanisms may include protein truncation, loss of regulatory interactions, and partial reductions in activity, to name but a few. Yet, beside the raw fitness effect of mutations in the ancestral strains, other mechanisms not related to mitonuclear interactions may account for the observed patterns of mitonuclear and carbon source specificity. Fitness of the ancestral strains (**Fig 1B**, **Fig S2**) differs markedly between mitochondrial backgrounds. If the fittest backgrounds are indeed approaching fitness optimum, as hypothesized above, and given patterns of diminishing returns epistasis common in experimental evolution (Chou et al. 2011; Wiser et al. 2013; Couce and Tenaillon 2015), the magnitude of fitness gains accessible to different mitonuclear pairs likely differed, constraining evolutionary paths to adaptation. Considering that initial fitness itself is influenced by mitonuclear interactions, this effect may prove difficult to deconvolute from other similarly influenced mechanisms. However, we did identify loci uniquely or preferentially mutated in mitonuclear backgrounds that otherwise display comparable initial fitness to closely related backgrounds (**Fig S8**). We also counted the number of non-synonymous mutation calls in evolved individuals. We found that the median number of mutations detected per evolved individual is two (**Data S3**). Hence, in evolved individuals, loss-of-function alleles normally interact with other mutant loci, contrary to our simplified experiment, performed in the ancestral backgrounds. This aspect appears critical when considering mutational order: selection for mutation at any given locus is contingent on the identity of mutations most likely to fix earlier during evolution (Weinreich et al. 2006). This itself may be influenced by mitonuclear interactions and illustrates the difficulty of predicting patterns of mutational specificity and evolutionary convergence based solely on fitness effects measured in the ancestral strains.

We also found that the nucleotide-level mutational profile observed in backgrounds NN and YY differed slightly but significantly from that of all other strains (**Fig S14**). This mutational bias would support the hypothesis that patterns of background-dependent evolutionary convergence may, at least in part, be dictated by differential access to beneficial mutations rather than mitonuclear interactions. Yet, because the nuclear genome, by virtue of its much larger size, chiefly determines mutational bias, this hypothesis does not account for differences in mutational specificity observed between backgrounds that carry the same nuclear genome (as illustrated in **Fig S8**). Moreover, fitness effects inferred from association tests in evolved mutants also correlated poorly with both convergent evolution signals and competition data (**Fig S13**). Together, these results indicate that different mitonuclear backgrounds may have access to different mutational spectra, in presence of multiple and complex epistatic networks. Mitonuclear interactions are a critical component of these epistatic networks and appear to exert a decisive impact on the path of evolutionary adaptation in *S. cerevisiae*.

## Discussion

The current model of mitonuclear evolution proposes that the mitochondrial genome, through interactions with nucleus-encoded genetic material, exerts an influence on evolutionary divergence of populations and eventually on the process of speciation. An elevated rate of evolution observed in the mitogenome can exert a pressure for compensatory evolution of the nuclear genome, leading to negative interactions between mitonuclear pairs that have evolved independently (Dowling 2014). Evidence of mitonuclear incompatibility between both closely related species (Niki et al. 1989; Osuský et al. 1997; Dey et al. 2000a; McKenzie and Trounce 2000; Špírek et al. 2000; Yamaoka et al. 2000; Kenyon and Moraes 2002; Lee et al. 2008; Chou et al. 2010; Ma et al. 2016) and geographically isolated populations of the same species (Paliwal et al. 2014; Wolters et al. 2015; Barreto et al. 2018; Wolters et al. 2018; Nguyen et al. 2020) corroborates this model. Yet, direct experimental evidence demonstrating the impact of mitonuclear interactions on evolutionary trajectories and outcomes was still lacking. We designed an experimental evolution study aimed at filling this gap. We mimicked the potential pressure exerted by the mitochondrial genome on the nuclear genome by swapping mitochondrial genotypes among yeast strains. We propagated these strains in a highly replicated manner for several hundred generations in conditions that either explicitly select for a mitochondrial function or relax the selective pressure exerted on this function. If the mitochondrial genome exerts a pressure on the nuclear genome, independent populations of the same nuclear genotypes but with distinct mitochondrial haplotypes should converge on different adaptive mutations.

### Mitonuclear interactions and their impact on evolutionary trajectories

The outcomes of our evolution experiment and the analyses herein support our prediction that specific mitonuclear interactions led to distinct evolutionary trajectories. Our phenotypic data shows the influence exerted on fitness of mitonuclear interactions in both ancestral and evolved individuals. Such effects depended on the conditions in which they were tested, the evolution regimen, and fitness proxies retained for analysis. Fitness in the evolved individuals that we isolated is on average comparable albeit not identical among the mitonuclear backgrounds under study. Genomic scrutiny in those individuals provides additional evidence for the impact of mitonuclear interactions on evolutionary outcomes. Thanks to our highly replicated dataset, we were able to detect signals of evolutionary convergence within starting genotypes at the whole-chromosome and gene levels, as observed in previous evolution experiments (Tenaillon et al. 2012). This allowed us to identify features of the yeast genome specifically or preferentially targeted by selection in a mitonuclear background-specific manner. This included changes that were specific to one single mitonuclear background, to the exclusion of closely related ones. We similarly detected environment-specific patterns of selection. Furthermore, recapitulating loss-of-function alleles at convergent loci in the ancestral strains of our experiment, we demonstrated that their effect on fitness, while generally favourable, varied markedly as a function of mitonuclear background and carbon source. Taken together, this evidence therefore upholds standing models of mitonuclear evolution, whereas specific interactions between the nuclear and mitochondrial genomes lead to divergent evolutionary trajectories.

Screening our collection of mitonuclear hybrids, we observed that the starting nuclear genome exerted a larger impact on fitness than did the mitochondrial genome. Highly replicated assessment of fitness in the ancestral strains confirmed this observation but indicated a modulating influence for the mitochondrial genome and mitonuclear interactions. Further supporting this view, we found evidence that the disruption of co-evolved mitonuclear pairs via cytoduction can impact the activity of mitochondrial enzymes and is associated with changes in mtDNA content. These founding contingencies in the experimental evolution represent, on their own, strong predictors of the divergent evolutionary trajectories predicted from different mitonuclear backgrounds. Indeed, the contingent evolutionary history of an organism creates a unique ensemble of constraints on the set of alleles likely favored by selection. In other words, it potentiates specific evolutionary pathways, and examples of this are reported in the experimental evolution literature (Blount et al. 2008).

Evolved individuals displayed greater phenotypic uniformity than their unevolved ancestors. Less adapted mitonuclear backgrounds readily caught up to fitter ones. This is coherent with the rapid and reproducible gains in fitness generally observed in the early stages of experimental evolution (Wiser et al. 2013) and with patterns of diminishing returns epistasis expected in fitter individuals (Chou et al. 2011; Couce and Tenaillon 2015). Evolved individuals display phenotypic uniformity in the environment in which they evolved, but also in conditions to which they were not explicitly adapted. This could be a sign of positive pleiotropy, meaning that many of the mutations selected in one environment tend to be adaptive in the other. More prosaically, we note that the two environments differ solely by the identity of the available carbon source. Hence, we posit that both environments are more alike than different, and alleles selected in either will tend to be adaptive in both by virtue of the same mechanisms. Despite this, we did observe several environment-specific mutant annotations. We also observed slightly lower fitness in fermentable medium for individuals evolved in non-fermentable medium. Therefore, environment-specific adaptive strategies still arose, which may steer the path of evolution towards more strongly distinct pathways over a longer evolutionary period.

Mitonuclear cooperation posits a complex network of genetic and physical interactions. Compatible with this framework, patterns of mitonuclear-specific evolutionary convergence and mitonuclear interactions presented in this study suggest selection affected by high-order epistasis. Recapitulating some of the selected mutations as loss-of-function alleles further argued for that view, as their effect in the ancestral backgrounds was dependent on the mitonuclear background. Furthermore, the locus preferences we could infer from the fitness of these reconstituted mutants did not match predictions from patterns of evolutionary convergence. Since we know that each evolved individual carries more than one mutation, we can reasonably posit that the effect of mutation at any locus in our mutants, and the probability for their selection during evolution, is affected by other mutations found in the same cells, especially those that arose earlier. The factor that unifies all these considerations is the founding difference in mitonuclear background.

### Limitations of the study

Our evolution experiment lasted approximately 300 generations. While published experimental evolution studies of a few hundred generations are common, this remains short on the evolutionary time scale. For comparison, this would correspond to roughly 7500 years of human evolution. While most studies cannot realistically last for tens of thousands of generations, a few thousand generations are commonplace (Lenski 2017). The typical workaround for short-term experimental evolution is increased replication (Mcdonald 2019). With more than 1300 starting populations, our experiment certainly meets the high replication standard, yet studies with similar levels of replication have reported lengths of up to a thousand generations (Lang et al. 2011; Lang et al. 2013). Despite this limitation, we were able to identify numerous and clear patterns of evolutionary convergence within genotypes. In addition, we could assign both mitonuclear and carbon source specificity to many mutant loci. Furthermore, it is uncertain that prolonged evolution would have increased the statistical power of our study. While it may have exacerbated patterns of specificity at some loci, over long periods, the accumulation of fixation events coupled to clonal interference and negative epistasis could have reduced the diversity of detectable patterns. However, it could also have enabled the detection of patterns that could only emerge after the fixation of earlier mutations. In short, by affording time for divergence, a longer evolution may have helped observe more sharply defined evolutionary trajectories for each mitonuclear background.

By evolving yeast in two environments that differed solely by the identity of the available carbon source, we hoped to test for the effect of explicitly selecting for a mitochondrial function, while minimizing confounding factors. In retrospect, comparing outcomes of evolution in two highly similar environments may have led to excessively similar outcomes, especially at the phenotypic level. Variation of an additional factor, such as nitrogen source, temperature, or even carbon source concentration, may have helped further differentiate populations evolving in distinct conditions, and better identify effects specific to mitochondrial function. Nevertheless, our experimental setup proved sufficient to identify carbon source-specific mutational patterns and compare patterns of divergence associated with different selective pressures.

The seven mitonuclear backgrounds that we submitted to experimental evolution were derived from just three original strains. We screened our collection of cytoductants for mitonuclear mismatches with readily detectable phenotypic differences, maximizing our chances of observing bona fide examples of mitonuclear interactions. This may have unduly restricted the breadth of observable patterns of mitonuclear interaction and evolution, biasing our results towards specific evolutionary outcomes. An alternative would have been to propagate a much wider diversity of mitonuclear backgrounds, potentially evolving our entire collection. This agnostic approach would have limited potential artefactual biases induced by the experimental design, at the expense of per background replication.

Most of the evidence presented in this study relies directly or indirectly on the results of sequencing. We did phenotype each of our evolved isolates. We also attempted recapitulation of some of the loss-of-function alleles identified by sequencing, generating corresponding knockouts in ancestral backgrounds. Yet, these experiments either imperfectly mimicked the conditions of evolution, were limited in scope or relied on simplifying assumptions. Indeed, systematic assessment of growth kinetics in evolved isolates was performed on solid medium for ease of replication and automation, whereas evolution was performed in liquid to accelerate evolution. Thus, phenotypic insight from growth on solid may not accurately reflect growth in the conditions of evolution. In contrast, high-sensitivity competition assays on recapitulated loss-of-function alleles assessed fitness effects in the exact conditions used for evolution. Yet, those assays concerned but a limited number of loci. For technical reasons, mutations to the mitochondrial genomes and aneuploidies had to be excluded from these tests. More generally, our restricted choice of loci may not reflect most mutations, biasing our view of fitness effects. Potentially more impactful is our simplifying assumption that mutations at the chosen loci were pure loss-of-function alleles, which we chose to recapitulate using whole gene knockouts. Careful reconstitution of the exact mutations identified by sequencing, while less scalable, may have provided a more accurate view of fitness effects. Finally, knockouts were prepared in ancestral backgrounds, ignoring epistatic effects incurred by other mutations accumulated in evolved individuals. We thus acknowledge that our phenotypic data represents an approximation, potentially complemented by more targeted future studies.

### Future directions

The model of mitonuclear evolution tested in this study predicts that evolutionary divergence is driven by the high rate of evolution experienced by the mitochondrial genome. In turn, the nuclear genome is believed to accumulate compensatory mutations aimed at maintaining compatibility. Data presented in this study support this model by showing the influence of specific mitonuclear interactions on the evolutionary trajectories and range of selected mutations observed in adapting mitonuclear hybrids. We also showed evidence of mitonuclear incompatibility and metabolic disruption in the mitonuclear hybrid ancestors of the experiment. In addition, we provided evidence for the resolution of the incompatibilities. However, this study shows no evidence about the site of compensatory evolution in either the nuclear or mitochondrial genomes. Strains evolved during our experiment may yet enable such demonstrations. Using cytoduction, we may generate novel mitonuclear hybrids by introducing an ancestral mtDNA or nuclear genome into the evolved strains. From a combination of growth, competition and enzymatic assays, along other cellular properties, we should identify which of the nuclear or mitochondrial genome is responsible for compensation. As an example, crossing evolved ND individuals with their ancestor, we could determine if evolved N nuclear genomes converged towards D, or if rather the D mitochondrial genome converged towards N. This experiment would provide an additional test for the standing model of mitonuclear evolution, potentially providing experimental support.

Our sequencing efforts have identified several mutant loci, with potential relevance to mitochondrial biology and mitonuclear evolution (see **Text S1**). Detailed scrutiny of the cell and molecular biology of adaptive mutations at these loci should provide insight into their underlying mechanisms, shedding light not only on paths of mitonuclear compensation, but also fundamental cellular processes. For example, aneuploidies at chromosome IV have been associated with increased gene dosage at *TSA2* in response to oxidative stress (Linder et al. 2017). We could therefore study the impact of overexpression of this gene in various mitonuclear backgrounds, testing for differential effects. Similarly, artificial expansion of the rDNA locus, as in (Kwan et al. 2016), may help investigate the adaptive influence of chromosome XII copy number on mitonuclear homeostasis. Long read sequencing at *HPF1* and *MNN4* loci in individuals mutant would provide an accurate map of the associated CNVs. This would enable their reconstitution in various backgrounds. Along with already available *RIM20* knockouts, expression of altered *HPF1* and *MNN4* constructs could help to study cell wall mannoproteins. We could further study their role in yeast buoyancy, in addition to their impact on respiration, oxidative stress and chronological lifespan.

### Summary and conclusion

In this study, we used experimental evolution of yeast mitonuclear hybrids and whole-genome sequencing of the resulting individuals to uncover patterns of evolutionary convergence specific to unique mitonuclear backgrounds. We also confirmed that phenotypic outcomes of evolution are primarily contingent on the nuclear background, with a modulating influence for mitonuclear interactions. We further showed that mitonuclear interactions exert an influence on the molecular mechanisms of adaptation, potentially constraining and potentiating long term evolution. To our knowledge, this study represents the first direct experimental demonstration of mitonuclear interactions inflecting the evolutionary trajectories of adapting populations. In addition to the support provided to standing evolutionary theory, the strains generated in this study and the accumulated data provide material for further studies of mitonuclear evolution and the cell biology of yeast.

## Materials and Methods

### Culture media

Below are the recipes for the media used in this study. Medium ingredients were purchased from BioShop (Burlington, ON, Canada). Experimental evolution, with its starter cultures, as well as growth and competition assays were all performed in media assembled from ingredients of the same production lot (see below).

#### Yeast peptone dextrose (YPD)

10 g/L yeast extract, 20 g/L tryptone, 20 g/L glucose. 2XYPD is assembled by doubling the concentration of all components in YPD.

#### Yeast peptone glycerol (YPG)

10 g/L yeast extract, 20 g/L tryptone, 20 g/L glycerol.

#### Yeast peptone ethanol glycerol (YPEG)

10 g/L yeast extract, 20 g/L tryptone, 30 g/L ethanol, 20 g/L glycerol.

#### Enhanced YPD (eYPD)

15 g/L yeast extract, 40 g/L tryptone, 40 g/L glucose, 5 g/L malt extract, 2.5 g/L glycerol, 0.4 g/L L-cysteine hydrochloride, 0.83 g/L K_2_HPO_4_, 3 g/L Tris pH 6.0.

#### Amino acid dropout without uracil and without histidine

The following compounds were mixed in powder form: 0.5 g L-adenine sulfate dihydrate, 2 g L-arginine hydrochloride, 2 g L-aspartic acid, 2 g L-glutamate monosodium salt, 10 g L-leucine, 2 g L-lysine monohydrochloride, 2 g L-methionine, 2 g L-phenylalanine, 2 g L-serine, 2 g L-threonine, 2 g L-tryptophan, 2 g L-tyrosine, 2 g L-valine.

#### Enhanced synthetic complete (eSC)

3.48 g/L yeast nitrogen base without ammonium sulfate and without amino acids, 2 g/L monosodium glutamate, 40 g/L glucose, 2.5 g/L glycerol, 0.4 g/L L-cysteine hydrochloride, 2.4 g/L amino acid dropout without uracil and without histidine, 0.83 g/L K_2_HPO_4_, 3 g/L Tris pH 6.0.

Solid medium was made by adding 1.5%-2% agar to the above recipes.

### Screen of collection of yeast cybrids

A collection of yeast cybrids was previously constituted, as described in (Paliwal et al., 2014; Wolters et al., 2018). It contains all possible combinations of nuclear and mitochondrial genomes from a set of fifteen strains of *S. cerevisiae*, in four biological replicates. Each strain in the collection was assayed for growth in non-fermentable YPG medium as follows. All manipulations were performed following standard sterile technique. Five microliters from thawed glycerol stocks of the collection were transferred to YPD evolution plates and incubated for 48 hours at 30°C, 65% relative humidity (RH), with shaking at 165 rpm in a microplate incubator shaker (Multitron, Infors HT, Basel, Switzerland). With the help of a liquid handling robot (Freedom Evo, Tecan, Männedorf, Switzerland), 5 µL from these prolonged cultures were transferred to the wells of a flat bottom polystyrene microtiter plate (Greiner, Kremsmünster, Austria), each well containing 235 µL of YPG and a 2.5 mm diameter glass bead. This second plate was covered with a transparent lid, with sides wrapped in three layers of parafilm (Bemis, Sheboygan Falls, WI, USA). Plate was placed in a Tecan Infinite M Nano microplate reader and incubated for 48 hours at 30°C, recording A_595_ at 20 min intervals. Each 20 min interval consisted of 13 min orbital shaking (6 mm diameter), 5 min absorbance measurement, and 2 min resting time. Raw growth curve data was fitted to a modified Gompertz model, as in (Zwietering et al., 1990). Area under the fitted curve was calculated using the trapezoidal rule.

### Evolution plates

Experimental evolution and related experiments were performed in standardized microtiter culture vessels, assembled as follows. Flat-bottom polypropylene microtiter plates, in 96 well format (cat. nb. 951040005, Eppendorf, Hamburg, Germany) served as reusable, autoclavable culture vessels. A clean 2.5 mm diameter glass bead (Biospec, Bartlesville, OK, USA) was placed in each well. Wells were filled with 240 µL media, then sealed with an adhesive foil seal (VWR, Wayne PA, USA). Plates were next covered with clean polypropylene lids held with autoclave tape. Plates were sterilized with a 20 min liquid autoclave cycle at 121°C. Plates were left at room temperature to cool and dry. Plates were then wiped of any residual water, placed in airtight plastic bags, and stored at 4°C until use. Before use, plates were equilibrated at room temperature and centrifuged at 2000 rpm for 2 min. Immediately before inoculation, lids were removed, and foils seals discarded. Following inoculation, plates were sealed with breathable adhesive membranes (VWR, Wayne PA, USA), and covered with polypropylene lids held with masking tape at all four corners. For re-use, plates were emptied, collecting and rinsing beads. Beads were sanitized by a 1 hr+ incubation in warm 10% bleach, rinsed thoroughly with tap water, and dried at 56°C in a Pasteur oven. Plates were washed in soapy water, rinsed thoroughly with tap water, and left inverted at room temperature until fully dry.

### Assessing the growth of ancestral and evolved individuals on fermentable and non-fermentable solid media

High-throughput assessment of growth in evolved individuals and their ancestral strains was performed by recording the growth of colonies on YPD and YPG agar medium. Replication and array of samples in preparation for growth curves were performed with robotically manipulated pin tools (BM5-SC1, S&P Robotics, Toronto ON, Canada). Thawed glycerol stocks were replicated from 96-well plates to omnitrays (Nunc, Rochester, NY, USA) in 96-array format on YPD agar medium. Replicas were incubated at 30°C until large, well-defined colonies were visible. Replicas were then cherry-picked and re-arrayed to 384-array format on YPD agar, randomizing positions. Following outgrowth, plates were combined into 1,536-array format on YPD agar omnitrays. Evolved individuals of a given mitonuclear background, evolved in both fermentable and non-fermentable conditions, were printed on a single 1,536 array. Each evolved individual was printed as six replicates. Hence, one 1,536 array was prepared per mitonuclear background for a total of seven arrays, whereas nine replicates of each of the ancestral strains were printed onto all seven arrays. To avoid border effects, top and bottom two rows of the arrays were printed with a filler strain, as were the two extreme left and right columns, and these were excluded from analysis. Next, these randomized arrays in 1,536 format were outgrown, then replicated on YPD and YPG agar. The replicas were incubated at 30°C for 48 hrs (YPD) or 72 hrs (YPG) in a spImager custom robotic platform (S&P Robotics), recording photographs of the plates at 2-hour intervals. The outgrown replicas were replicated to fresh plates for two additional rounds of growth. Data from the second and third rounds was retained for analysis.

### Analysis and quantitation of growth on solid medium

Colony sizes from pictures were measured with R package Gitter (Wagih and Parts, 2014). Growth curves were drawn for each colony, and fitted to a modified Gompertz model, as described above, extracting estimates of the maximum specific growth rate (µ) and carrying capacity (A). Ancestral controls were used to estimate plate-specific effects on growth parameters and to normalize estimates across arrays. Specifically, the effect of a given plate on a growth parameter was calculated by dividing each individual ancestral estimate by the geometric mean of the ancestral estimates over all plates in each carbon source, then extracting the median of that ratio over all ancestral estimates from the same plate. The normalization factor thus obtained was used to multiply all estimates from a plate. True value for growth parameters of any given strain was estimated using the median (if n ≥ 25) or geometric mean (if n < 25) over all replicates, with confidence intervals obtained by bootstrapping. A series of two-way ANOVAs with interaction was performed on growth data with nuclear and mitochondrial backgrounds as factors, Separate ANOVAs were conducted for each combination of fitness proxy, evolution regimen, and carbon source.

### Preparation of whole extracts from strains of *S. cerevisiae* for enzyme assays

Precultures were inoculated by transferring 5 µL from thawed glycerol stocks to 5 mL 2XYPD medium. Cultures were incubated with shaking at 30°C for 24 hrs. Half a milliliter from precultures was used to inoculate 5 mL YPG, and this new culture was incubated with shaking at 30°C for 24 hrs. This second culture was diluted in 45 mL of pre-warmed YPG medium. This subculture was incubated at 30°C with shaking until it reached A_600_=0.5-1.0. Cells were pelleted by centrifugation, then suspended in water. Cells were pelleted once again by centrifugation, washed in 25 mM potassium phosphate pH 8.0, then frozen as pellets at -80°C. After overnight freezing, pellets were thawed on ice, and suspended in 0.5 mL of freshly prepared, ice-cold 25 mM Tris-HCl pH 7.5, 1 mM EDTA, 100 mM NaCl + cOmplete Mini, EDTA-free Protease Inhibitor Cocktail (Roche, Basel, Switzerland). Suspension was bead beaten at 4°C for 10 cycles of 2 min beating, 2 min rest on ice. Lysate was frozen at -80°C.

### Enzyme assays on whole cell extracts from *S. cerevisiae*

Enzyme activities were determined spectrophotometrically using a Mithras LB940 microplate reader (Berthold technologies, Germany) and data analyzed with MikroWin 2010 software (Labsis Laborsysteme, Germany). Enzymatic capacities were expressed as mU•mg proteins^−1^ (U•mg proteins^−1^ in the case of CAT), where U refers to 1 µmol of substrate transformed to product per minute. Chemicals were purchased from Sigma-Aldrich (Oakville, Ontario, Canada). Enzymatic assays were performed at 30°C in the following conditions:

#### Malate dehydrogenase (MDH, EC 1.1.1.37)

MDH activity was measured at 340 nm for 4 min, following the oxidation of NADH (ε_340_ = 6.22 mM^−1^•cm−^1^). The medium consisted of 100 mM potassium phosphate, 0.2 mM NADH and 0.5 mM oxaloacetate (OAA), pH 7.5. The background activity in absence of sample was subtracted from the main results (Bergmeyer, 1983).

#### Mitochondrial complex I + III (ETS, EC 7.1.1.2 and 7.1.1.8)

ETS activity was measured at 490 nm using NADH as electron donor, following the reduction of p-iodonitrotetrazolium violet (INT, ε_490_ = 15.9 mM^−1^•cm^−1^) for 6 min. The medium consisted of 50 mM potassium phosphate, 0.85 mM NADH, 2 mM INT and 0.03% (v/v) triton X-100, pH 7.5. The specificity of the reaction was verified in absence of NADH and the residual activity was subtracted from the main results (Hunter-Manseau et al., 2019).

#### Cytochrome c oxidase (CCO, EC 7.1.1.9)

CCO activity was measured at 550 nm following the oxidation of cytochrome c (cyt c, ε_550_ = 18.5 mM^−1^•cm^−1^) for 6 min. Cyt *c* was reduced through the addition of 4.5 mM dithionite. The medium consisted of 50 mM potassium phosphate, 50 µM cyt c, 1 mM ADP and 0.03% (v/v) triton X-100, pH 7.0. The specificity of the reaction was verified in presence of 40 mM sodium azide (CCO-inhibitor), and the residual activity was subtracted from the main results (Lemaire and Dujardin, 2008; Spinazzi et al., 2012; Hunter-Manseau et al., 2019).

#### ATP-synthase (ATPase, EC 7.1.2.2)

ATPase activity (ATP hydrolysis) was measured at 340 nm following the oxidation of NADH (ε_340_ = 6.22 mM^−1^•cm^−1^) for 4 min. The medium consisted of 250 mM sucrose, 20 mM HEPES, 5 mM MgSO_4_, 0.35 mM NADH, 2.5 mM phosphoenolpyruvate (PEP), 2.5 mM ATP, 5 μM antimycin A, 4 units • mL^−1^ lactate dehydrogenase (LDH) and 4 units • mL^−1^ pyruvate kinase (PK), pH 8.0. The specificity of the reaction was verified in presence of 25 μM oligomycin (ATPase-inhibitor), and the residual activity was subtracted from the main results (Barrientos et al., 2009; Haraux and Lombes, 2019; Rodriguez et al., 2020).

#### Catalase (CAT, EC 1.11.1.6)

CAT activity was measured at 240 nm following the disappearance of H_2_O_2_ (ε_240_ = 43.6 M^−1^•cm^−1^) for 1 min. The medium was composed of 100 mM potassium phosphate, 0.1% (v/v) triton X-100 and 60 mM H_2_O_2_, pH 7.5. The background activity in absence of sample was subtracted from the main results (Page et al., 2009; Pichaud et al., 2010; Orr and Sohal, 1992; Bettinazzi et al., 2021; Christen et al., 2020).

#### Citrate synthase (CS, EC 2.3.3.1)

CS activity was measured at 412 nm for 6 min, following the increase in absorbance due to the reaction between 5,5’-dithiobis-(2-nitrobenzoic acid) (DTNB) and CoA-SH to form TNB (ε_412_ = 14.15 mM^−1^•cm^−1^). The medium was composed of 100 mM Tris-HCl, 0.1 mM DTNB, 0.1 mM acetyl-CoA (AcCoA), 0.15 mM oxaloacetate (OAA), pH 8.0. The specificity of the reaction was verified in absence of OAA and the residual activity subtracted from the main results (Spinazzi et al. 2012; Hunter-Manseau et al. 2019).

#### Mitochondrial complex II (SDH, EC 1.3.5.1)

SDH activity was measured at 600 nm using succinate as electron donor and following the reduction of 2,6-dichloroindophenol (DCIP, extinction coefficient ε_600_ = 19.1 mM^−1^•cm^−1^) for 6 min. To fully activate the enzymatic complex, samples were incubated for 10 min at the assay temperature (30°C) in a reaction medium consisting of 50 mM potassium phosphate, 20 mM succinate and 5 mM MgCl_2_, pH 7.5. The reaction was started by addition of 50 µM DCIP, 65 µM ubiquinone1 (CoQ1), 4 µM antimycin A, 2 µM rotenone and 10 mM sodium azide. The specificity of the reaction was verified in absence of CoQ1 and the residual activity subtracted from the main results (Barrientos et al. 2009; Breton et al. 2009; Spinazzi et al. 2012; Hunter-Manseau et al. 2019).

#### Protein content

Protein concentration (mg•mL^−1^) was determined with the BCA assay (Smith et al., 1985).

### Preparation of starter cultures for experimental evolution

All procedures were performed under laminar flow within a sterile enclosure. This protocol was performed identically for strains of each of the following mitonuclear backgrounds: 273614N^273614N^ (NN), 273614N^DBVPG6044^ (ND), DBVPG6044^273614N^ (DN), DBVPG6044^DBVPG6044^ (DD), DBVPG6044^Y12^ (DY), Y12^DBVPG6044^ (YD), Y12^Y12^ (YY). Streaks from each of the four biological replicates from each of the mitonuclear backgrounds were prepared on YPEG agar, and incubated at 30°C for 48 hrs. From each streak, 24 colonies were picked at random with sterile micropipette tips and inoculated into YPD evolution plates. Hence, 96 cultures were launched from each of the seven mitonuclear backgrounds submitted to experimental evolution. Plates were incubated overnight at 30°C, 65% RH with shaking at 200 rpm. Glycerol stocks were prepared by mixing 100 µL from the starter cultures with 100 µL 75% glycerol in the wells of conical bottom polypropylene plates (Greiner, Kremsmünster, Austria). Remainder of the cultures were used immediately to launch experimental evolution.

### Experimental evolution on mitonuclear hybrids of *S. cerevisiae*

Our collection of yeast cybrids contains four biological replicates of each mitonuclear background, each derived from a separate cybridization event. Twenty-four evolutions were founded from each of these cybrids, hence 96 replicate populations of each of seven mitonuclear backgrounds were propagated by daily serial passage in both fermentable (YPD) and non-fermentable (YPG) medium for 58 days. All procedures were performed under laminar flow within a sterile enclosure. Starter cultures were used to inoculate a first batch of experimental evolution cultures. With the help of a liquid handling robot (Freedom Evo 150, Tecan, Männedorf, Switzerland), 5 µL from each starter culture were transferred to YPD evolution plates, and in parallel to YPG evolution plates. Tips used for transfer were washed with sterile water and kept for further transfer of the same cultures. Plates were incubated for 24 hrs at 30°C, 65% RH with shaking at 200 rpm. Evolving populations were then transferred to new evolution plates for 58 consecutive days. On day 2 and every 7 days afterwards until day 58, glycerol stocks were prepared by mixing 100 µL from the evolving populations with 100 µL 75% glycerol in the wells of conical bottom polypropylene plates.

### Isolation of independently evolved individuals

Glycerol stocks from day 58 of the evolution experiment were thawed at room temperature. Ten microliters from each stock were transferred to the surface of a YPD agar petri dish. The resulting drop from the glycerol stock was streaked for single colonies. Petri dishes were incubated for 48 hrs at 30°C. From each Petri dish, a single colony was picked at random using a sterile tip and placed into a YPG evolution plate. Plates were incubated for 24 hrs at 30°C, 65% RH with shaking at 200 rpm. Glycerol stocks were prepared by mixing 100 µL from the starter cultures with 100 µL 75% glycerol in the wells of conical bottom polypropylene plates (Greiner, Kremsmünster, Austria).

### Preparation of total DNA from evolved individuals and ancestral strains

Total DNA from each individual evolved strain and all ancestral strains was extracted and purified as starting material for the preparation of next-generation sequencing libraries. A custom protocol using the DNeasy Blood and Tissue Kit (Qiagen, Venlo, Netherlands) was applied in 96-well format. Each strain was inoculated from glycerol stock in 4 × 1 mL YPD medium, in a polypropylene 96-well deep well plate (DWP). Each DWP was covered with a breathable adhesive membrane and incubated for 24 hours at 30°C with shaking at 200 rpm. Cells were harvested by centrifugation for 10 min at 5000 x g and suspended in 250 µL water. Suspensions of each strain were pooled into a single well, and centrifuged once more for 10 min at 5000 x g. Pellets were suspended in 600 µL of 1 M sorbitol, 100 mM sodium EDTA, 14 mM β-mercaptoethanol, supplemented with 6 U zymolyase (Bioshop, Burlington ON, Canada), then transferred to collection microtubes provided with the DNeasy kit, and incubated at 30°C for at least 30 min. Resulting spheroplasts were pelleted by centrifugation for 10 min at 5000 x g, suspended in 180 μl Buffer ATL + 20 μl proteinase K solution, and mixed thoroughly. Lysis suspensions were incubated at 56°C for at least 30 min, mixing occasionally to disperse the samples. Samples were shaken vigorously for 15 s, then centrifuged briefly to collect all liquid at the bottom of tubes. Samples were supplemented with 4 μL of 100 mg/mL RNase A, mixed vigorously, and incubated at room temperature for at least 5 min. A 50:50 mix of buffer AL and ethanol was prepared fresh, and 400 μL were dispensed to each tube, mixing once again by vortexing. Lysates (max 900 μL) were applied to DNeasy 96 purification columns. Columns were centrifuged for 10 min at 3800 x g. Columns were washed with 500 μL buffer AW1, centrifuging for 5 min at 3800 x g. Columns were further washed with 500 μL buffer AW2, centrifuging for 15 min at 3800 x g. Columns were next placed on a clean rack of elution microtubes and 100 μL of buffer AE were added to each sample. Samples were incubated for at least 1 min at room temperature, then centrifuged for 2 min at 3800 x g. Each eluate was next supplemented with 4 μL of 10 mg/mL RNase A, agitated vigorously, and incubated at room temperature for 15 min. The RNase-treated eluates were further purified by mixing 50 μL from each sample with 20 μL beads and incubating at room temperature for 5 min. After the bead mixture was applied to a magnet for 60s, liquid was removed from the beads, which were washed twice with 200 μL of 80% ethanol. Beads were suspended in 40 μL of 10 mM Tris-HCl pH 8.0 and incubated for 5 min at room temperature. Eluate was separated from beads using a magnet, as described above. Sample quality was evaluated by agarose gel electrophoresis. Purity and concentration of DNA was assessed by measuring absorbance at 260 nm and 280 nm using a NanoDrop spectrophotometer (ThermoFisher Scientific, Waltham MA, USA). Concentration of DNA in the sample was further estimated using the AccuClear Ultra High Sensitivity dsDNA Quantitation Kit (Biotium, Fremont CA, USA) following manufacturer’s instructions.

### Preparation of next-generation sequencing libraries

Sequencing libraries were prepared using the Tagment DNA Enzyme and Buffer Large Kit (Illumina, San Diego CA, USA), the KAPA HiFi HotStart ReadyMix 2X (Roche, Basel, Switzerland) and custom DNA barcodes. Genomic DNA samples were diluted with water to a target concentration of 2.5 ng/μL or less. Tagmentation reactions were assembled by mixing 1.25 μL tagment DNA buffer, 0.25 μL tagment DNA enzyme and 1 μL gDNA dilution. Reactions were incubated at 55°C for 5 min. Custom DNA barcodes were added by PCR as follows. Tagmented DNA was mixed with 3.76 μL 2X KAPA ReadyMix, and 0.625 μL both primers. Primers had sequences 5’AATGATACGGCGACCACCGAGATCTACAC-(8-nucleotide custom i5 barcode)-TCGTCGGCAGCGTC3’ and 5’CAAGCAGAAGACGGCATACGAGAT-(8-nucleotide custom i7 barcode)-GTCTCGTGGGCTCGG3’. A collection of 32 custom i5 barcodes and 48 custom i7 barcodes was used, enabling the preparation of 1,536 unique barcode combinations. PCR was performed by cycling as follows: 72°C for 3 min, 98°C for 2:45 min, then 8 cycles of 98°C for 15s, 62°C for 30s, 72°C for 3 min, followed by a 1 min final extension at 72°C. Library preparation was completed by reconditioning PCR. Barcoding reactions were mixed with 8.5 μL 2X KAPA ReadyMix, 0.5 μL of 10 μM primer P1 (5’AATGATACGGCGACCACCGA3’) and 0.5 μL of 10 μM primer P2 (5’CAAGCAGAAGACGGCATACGA3’). Reactions were then cycled as follows: 95°C for 5 min, then 4 cycles of 98°C for 20s, 62°C for 20s, 72°C for 2 min, followed by a 2 min final extension at 72°C. Final PCR products were diluted with 14 μL PCR grade water, then bead purified as described above, except that only 18 μL beads were added to the PCR reactions. Library concentration was estimated using the AccuClear Ultra High Sensitivity dsDNA Quantitation Kit following manufacturer’s instructions. A subset of the libraries was inspected with a BioAnalyzer 2100 using a High Sensitivity DNA chip (Agilent Technologies, Santa Clara CA, USA) for insert size and monodispersity. Equimolar amounts of all libraries were pooled and sequenced on an Illumina NovaSeq6000 S4 PE150 sequencing lane at the Génome Québec Expertise and Service Center (Montréal QC, Canada).

### Pre-treatment of sequencing results and alignment to reference genomes

Quality control was performed individually on each sequencing library using FastQC (Andrews, 2010), and results were aggregated using MultiQC (Ewels et al., 2016). Sequencing adapters were trimmed with Trimmomatic (Bolger et al., 2014). Overlapping read pairs were merged using BBmerge (Bushnell et al., 2017). Single read files from the BBmerge output were concatenated into single files. Primary read mapping to reference genomes was performed using bwa mem (Li and Durbin, 2009). Libraries were mapped to the published genomes of their associated ancestral strains. The *S. cerevisiae* 273614N reference genome sequence was obtained from the diArk database (Hammesfahr et al., 2011) as published in (Liti et al., 2009). Reference genome sequences for *S. cerevisiae* strains DBVPG6044 and Y12 were obtained from NCBI BioProject PRJEB7245, BioSamples SAMEA2757762 and SAMEA2757763, respectively, as published in (Yue et al., 2017). Chimeric reference genomes were assembled for cybrid strains by manually replacing the mitochondrial genome contig published for the nuclear parent’s genome with that published for the mitochondrial parent. Alignment was performed separately for merged and paired reads. Hence, BAM outputs from bwa mem were merged and then sorted with samtools (Li et al., 2009). Following the addition of read groups and deduplication with picard (Broad Institute, 2016), indel realignment was performed using GATK3 (McKenna et al., 2010; van der Auwera and O’Connor, 2020). Alignment and depth metrics were collected using picard and samtools, respectively. Whole-genome pileups were extracted for each sequencing library with samtools.

### Estimating the fraction of mtDNA per cell

The fraction of sequence bases mapped to coding regions of the mitochondrial genome was used as a proxy for copy number, structural or other changes incurred to the organellar genome. This was computed from pileup, summing the depth of coverage with base quality greater than 20 in known coding regions of the reference mitochondrial genomes. This sum across the mitochondrial genome divided by the same sum over the whole genome provided an estimate of the fraction of coding mitochondrial DNA in each cell.

### Identification of aneuploidies from sequencing data

Aneuploidies were called from pileup. Telomeric and subtelomeric regions were excluded from this analysis by masking 25 kb at both ends of each chromosome. High quality depth of coverage was computed at each position by counting reads with both base and mapping quality above 30 on the phred scale. Finally, any position found with average depth of coverage in all biological replicates of the ancestral strains above 5 times the median depth over the whole genome was also excluded from analysis. Next, the ratio between the mean depth of the chromosome and that of the rest of the genome was calculated. If this ratio was between 0.75 and 1.25, a p-value for the mean coverage of the chromosome was computed, assuming normal distribution of depth of coverage across the genome, with mean and standard deviation computed over the rest of the genome. If this p-value was smaller than the length of the chromosome divided by the length of the whole genome, an aneuploidy was suspected. Aneuploidy candidates were confirmed visually by inspecting heatmaps of depth of coverage along the length of chromosomes.

### Primary identification of copy number variants (CNVs) from sequencing data

Primary CNV calling was performed essentially according to (Yoon et al., 2009). Analysis was performed from filtered pileups, as described above for aneuploidies. Each chromosome was divided in 100 bp windows, extracting mean depth of coverage for each window. Average and standard deviation on window means were computed across chromosomes, enabling computation of a z-score and p-value for each window. Next, whole chromosomes were scanned with rolling intervals of various window sizes, starting with 2 windows in length. If the maximum p-value for windows within an interval fell below a threshold, determined by both window and chromosome length, the interval was marked as a candidate CNV. Increasing interval sizes were used until a threshold determined by both interval and chromosome length was reached. Overlapping CNV candidates were merged. Any candidate with median depth between 0.75x and 1.25x the median chromosome depth was excluded. A p-value was computed for the mean depth of each candidate, based on mean and standard deviation over windows. If this p-value was greater than 10^−6^, the candidate was excluded. Finally, if the same area had mean depth of coverage between 0.75x and 1.25x the chromosome-wide median depth in the ancestral strains, the candidate CNV was also excluded.

### Resolution of artefactual CNV calls caused by misalignment at homologous loci

Reference genome sequence at CNV coordinates were blasted against the whole reference genome sequence, identifying loci with high homology to CNV candidates. Blast hits with an E-value of less than 1.0, length above 150 bp and percent identity with the CNV locus above 80% were retained for further scrutiny. Sequences at the CNV call and homologous hit loci were aligned with MUSCLE (Edgar, 2004), identifying locus specific positions. A system of linear equations was built from the sums of each base calls at each locus specific positions in the alignment. Multiple linear regression was then used to identify the most likely depth of coverage, with 95% confidence interval, at each homologous locus. If the inferred depth was between 0.75x and 1.25x the chromosome median depth, and the confidence interval on depth at the homologous locus did not overlap with the confidence interval for depth over the whole chromosome, a CNV call was asserted. Otherwise, or if locus specific information was too scarce to lift ambiguity caused by homology, CNV call was not retained. Applying this methodology, several CNVs were rejected, others were retained, while a subgroup was reassigned to more likely loci.

### SNV calling

Primary SNV calling was performed using two parallel methods, starting with GATK HaplotypeCaller. SNV calls were combined into a single GVCF file using GATK CombineGVCF. Initial filtering was performed with GATK FilterVariants and SelectVariants. Additional filtering was done with vcftools. A list of SNVs was produced in csv format using bcftools.

In parallel, custom SNV calling was attempted directly from pileup. The following regions of the genome were filtered out of pileups in preparation for SNV calling: high coverage regions in the ancestral strains, subtelomeric regions, positions with mean base and mapping quality below 30, positions with depth of coverage below 10, and positions with N or otherwise missing sequence in the reference. A baseline error model was built, identifying the number of each base call expected at each position, considering the reference base and mean base quality at that position, and the average error rates observed in the data. A chi^2^ test for goodness of fit was performed, comparing the expected base counts predicted by the error model with the observed base counts. An E-value was obtained by multiplying this p-value with the number of positions in the genome. Positions with an E-value < 1.0 were retained as potential SNVs, while any position with non-reference base call count less than 25% of total depth was rejected.

The intersection between the results of both primary SNV calling pipelines was kept as the overall primary list of SNVs. SNV calls were additionally filtered for strand, base quality and mapping quality biases. A few libraries were observed to display SNVs that overlapped strongly with those from libraries that were found in close proximity on library preparation plates. Those SNVs were flagged as potential cross-contamination artifacts and removed from further scrutiny. Furthermore, SNVs found in two or more libraries that shared a barcode were tagged as potential index hopping artifacts and removed from SNV calls. Moreover, the same substitutions arising repeatedly at the exact same position were deemed unlikely: any SNV encountered in four libraries or more was rejected. SNVs expected to arise from alignment of reads to the wrong reference genome (e.g. DBVPG6044 reads mistakenly aligned to the Y12 reference genome) were also removed. Next, using Fisher’s exact test, SNV calls were identified that were more compatible with the homozygous non-mutant genotype than the heterozygous or homozygous mutant scenarios, and thus rejected. Finally, libraries that presented an unusually high number of SNVs (more than 300 SNVs and more than 2 standard deviations above average number of SNVs per library) were rejected. A heterozygous genotype was called for many SNVs, despite uniformly haploid starting populations. A small sample of evolved individuals were thus assessed for ploidy, as described above, showing diploidization in approximately half sampled isolates (**Fig S15**).

### Annotation of CNV and SNV calls

Annotations of the S288c reference genome (Engel et al., 2022) were used for annotating SNV and CNV calls in evolved isolates. Sequences from mutation-carrying loci were blasted against the S288c reference genome to identify corresponding coordinates. Annotations corresponding to these locations were identified with Yeastmine (Balakrishnan et al., 2012).

### Analysis of dissimilarity between mutational profiles

Mutational profiles were built for each combination of mitonuclear background and evolution regimen (evolutionary circumstances) by counting mutations associated with each mutant annotation. Bray-Curtis dissimilarity was computed between the mutational profiles of all pairs of evolutionary circumstances. Non-parametric multi-dimensional scaling was performed from the resulting dissimilarity matrix, preserving the best from 10 000 independent initializations of the algorithm. The algorithm was run for a maximum of 3000 iterations, declaring convergence using a stress tolerance threshold of 10^−9^.

### Specificity of mutant loci for carbon sources and nuclear and mitochondrial backgrounds

The same strategy was applied to test the specificity of mutant loci for carbon source, nuclear background, mitochondrial background, and all combinations thereof. For each locus, a contingency table was assembled reporting counts of mutations within and outside the locus in each category. This table was submitted to a chi^2^ test of independence, yielding a p-value. Mutual information (MI, reported as dits) was calculated from the same contingency table. Significance thresholds on chi^2^ p-values were corrected for false discovery rate on all loci. Loci with chi^2^ p-value below threshold and mutual information two standard deviations above mean MI were deemed non-randomly distributed between categories, and thus displaying specificity.

### Analysis of association between phenotypic changes and mutant loci

Quantitative phenotypic data, including fitness proxies (carrying capacity and growth rate) in both fermentable and non-fermentable media, as well as the fraction of mtDNA per cell, are available for each evolved mutant. A list of annotated mutations is also associated with each mutant, enabling tests of the statistical association between mutant loci and phenotypic changes. For each mutant locus and each phenotype in each carbon source, the association was tested as follows. A t-test was performed to determine if the fitness of individuals that display mutations at the locus of interest differed significantly from fitness of their ancestral strains, yielding a p-value. In parallel, using the same data, effect size of mutation at the locus was evaluated by calculating Cohen’s *d* corrected for sample size, with associated standard error. P-value of obtaining a given effect size at least as extreme by chance was computed empirically by drawing 10 000 random samples of the same size among mutants of the same mitonuclear background and calculating an effect size for each. These calculations were performed separately for each mitonuclear background since ancestral strains displayed widely differing phenotypes. P-values obtained for each mitonuclear background were combined using Stouffer’s method, with the square root of sample sizes used as weights. Significance thresholds were corrected for false discovery rate applying the method of Benjamini and Hochberg on all loci as described in (Kuo, 2017). Loci with both p-values below threshold were deemed significantly associated with quantitative changes in phenotype.

### Generation of fluorescent knock-out mutants at putative loss-of-function loci

First, GFP- and mCherry-tagged derivatives of all ancestral strains were generated from three biological replicates. Parental strains all carried the *MATa ho::hphMX ura3::KanMX* genotype. They were transformed with both the GFP and mCherry selection cassettes, which shared the same general structure. The expression of both fluorescent proteins was placed under the control of the *S. cerevisiae TEF2* promoter and *ENO1* terminator. Fluorescent protein expression cassettes were fused to the *natNT2* selection cassette. This full-length cassette was flanked on both extremities with 40 bp homology to the *HIS3* locus and transformed into the ancestral strains using the lithium acetate method. Transformants were selected on eYPD agar medium (as described in *Culture media* subsection above) supplemented with 100 μg/mL hygromycin, 100 μg/mL geneticin and 75 μg/mL nourseothricin. Integration of the fluorescent protein expression cassettes at the *HIS3* locus was confirmed by growth in presence of nourseothricin, loss of histidine prototrophy, detection of a fluorescent signal by flow cytometry and successful PCR amplification at the 5’ and 3’ junctions between the *HIS3* locus and heterologous sequences. Knockouts of each of the following genes were prepared in all fluorescent derivatives of the ancestral strains: *AMN1, GPB2, IRA2, MNN4, RIM20, SIR3, SIR4*, and *SUC2*. Open reading frames at each of these loci were deleted by transformation with the *Kluyveromyces lactis URA3* expression cassettes amplified from plasmid pUG72 (Gueldener et al., 2002) and flanked in 5’ and 3’ with 40 bp homology to the promoters and terminators of the target genes. Transformants were selected on eSC agar devoid of uracil. Knockouts were scored for uracil prototrophy; resistance to hygromycin, geneticin and nourseothricin; detection of a fluorescent signal by flow cytometry and successful PCR amplification at the 5’ and 3’ junctions between the target locus and heterologous deletion cassette.

### Competition assays between knockouts and their ancestral strains

Changes in fitness incurred by deletion at loci of interest were quantified by performing competition assays in fermentable and non-fermentable media, using a method inspired by (Breslow et al., 2008). Knockouts at each of the eight loci of interest, in the mitonuclear background under study, as three biological replicates, labeled with both GFP and mCherry were arrayed as glycerol stocks in 96-well format. Matching arrays of the fluorescently labeled ancestors were also prepared. Precultures in 96-well format of both the ancestral and knock-out strains were inoculated by transferring 5 μL from thawed glycerol stock arrays to YPD evolution plates. These preculture plates were incubated for 24 hrs at 30°C, 65% RH with shaking at 200 rpm. Outgrown precultures of the knockouts and matching ancestral strains were mixed by transferring 5 μL from both to YPD and YPG evolution plates. Control competitions of the mCherry and GFP labeled ancestral strains were also prepared, to account for differences in fitness caused by the fluorescent labels. For three consecutive days following mixing of the strains, competition cultures were propagated and transferred as described above for experimental evolution. On each day, outgrown cultures were diluted 1:50 in non-sterile water. Two thousand cells from each dilution were analyzed by flow cytometry, counting mCherry and GFP positive cells. Flow cytometry measurements were performed with the help of a Guava easyCyte HT (Luminex, Austin TX, USA) or Accuri C6 (BD, Franklin Lakes NJ, USA) instrument. All microbiological procedures were performed under laminar flow within a sterile enclosure, except for endpoint measurement by flow cytometry. All pipetting was performed with the help of a liquid handling robot (Freedom Evo, Tecan, Männedorf, Switzerland). Triplicate competitions were performed on both fluorescent derivatives of all biological replicates in both carbon sources.

### Analysis of competition data

The log_2_ ratio of mCherry count over GFP count was calculated for each time point in each competing culture. The log_2_ ratio at time zero was subtracted from time points of a given competition so that samples may share a common y-intercept. Linear regression was performed on all data points derived from the same biological replicate, fitting to the following model:

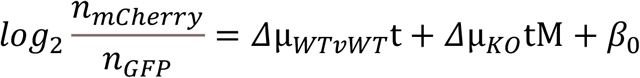

where n_mCherry_ is the number of mCherry-labeled cells, n_GFP_ is the number of GFP-labeled cells, *Δµ*_WTvWT_ is the difference in growth rate between the mCherry and GFP labeled ancestor, t is time, Δ*µ*_KO_ is the difference in growth rate between the knock-out and its ancestor, M is a variable with value 0 when a time point belongs to a ancestor vs ancestor competition, and 1 when it belongs to a knock-out vs ancestor competition, and β_0_ is a constant. The fit thus provides an estimate for the change in growth rate associated with knocking out the gene of interest in the current mitonuclear background. Standard error (SE) on this coefficient estimate is used to test whether the coefficient is null, and to compute a sum-of-squares error (SS) on the estimate. Estimates for three biological replicates and two fluorescent labels thus provided six estimates for each locus in each mitonuclear background. They were combined into a single estimate by calculating an average weighted by sample size. P-values were combined using Stouffer’s method, weighted by sample size. Similarly, sum-of-square errors were combined by summing them and adding the product of sample size with the square deviation of each estimate from the overall mean estimate. Mean fitness effect estimates, and the combined sum-of-squares were used to perform ANOVAs to test if fitness effect of a given locus in each carbon source differed between mitochondrial backgrounds, with accompanying Tukey HSD tests on pairwise differences between backgrounds. Similarly, ANOVAs were performed to determine if fitness effects of loci differed on given mitonuclear backgrounds. Two-way ANOVAs were performed to determine significance of nuclear, mitochondrial, and mitonuclear interaction effects on growth rate changes. Spearman rank correlation was calculated between fitness effects estimated by competition assays and frequency of mutations at the loci in each background, as determined from sequencing data.

### Mutational bias

We counted every possible single nucleotide change among all SNV calls in our dataset, and assigned them to their corresponding mitonuclear background. Counts of for each mitonuclear background were compared to the dataset-wide counts applying a chi^2^-test of independence, with a p-value threshold of 0.05.

### Assessment of ploidy in select strains

Ploidy was determined for all ancestral strains and a subset among evolved individuals, following a method inspired by (Charron et al. 2019). Thawed glycerol stocks of the strains were streaked for single colonies on YPD agar. Single colonies were picked in triplicate and inoculated into 1 mL YPD, in wells of a polypropylene DWP. DWPs were incubated for 24 hrs at room temperature. Cells were washed with 2 × 1 mL water, then fixed for 1 hr in 70% ethanol. Ethanol was removed by washing cells twice with 1 mL water. Cells were next incubated overnight at 37°C in 1 mL of freshly prepared 0.25 mg/mL RNase A. Cells were washed twice with 1 mL of 50 mM sodium citrate pH 7.0. A small volume (25 μL) of the washed cell suspension was diluted in 225 μL of 0.667 μM SYTOX green (Thermo Fisher Scientific) and incubated in the dark at room temperature for 1 hr. Cell density in the stained sample was adjusted to approximately 500 cells/μL, and analyzed by flow cytometry, recording 5000 events per sample. Mean green fluorescence was measured for cells in G1 phase. All manipulations were performed in triplicate.

## Acknowledgements

This work was supported by the Natural Science and Engineering Research Council of Canada (Discovery grant to CRL, RGPIN-2020-04844), the Fonds de recherche du Québec – Nature et Technologies (Team grant 2019-PR-254415 to CRL and SBr) and the Canada Research Chair in Cellular Systems and Synthetic Biology to CRL. DBP was supported by an award from the Fonds de recherche du Québec – Santé (FRQS). During part of the writing/reviewing, SBe was supported by the European Union’s Horizon 2020 research and innovation programme under the Marie Skłodowska-Curie grant agreement No 101030803. Raw sequencing data is available on the NCBI sequence read archive under BioProject accession number PRJNA878596.

## Authors contributions

Conceptualization: CRL, DBP, HLF, SBr

Data curation: DBP

Investigation: AKD, CB, DBP, SBe

Methodology: AKD, CRL, DBP, HLF, IGA, SBe, SBr,

Resources: CRL, HLF, SBr

Software: DBP

Supervision: CRL, HLF, SBr

Validation: AKD, DBP

Visualization: DBP

Writing – original draft preparation: DBP

Writing – review and editing: all authors

Formal analysis: DBP, SBe

Project administration: CRL

Funding acquisition: CRL and SBr

## References

Allen JF, Horner DS, Cavalier-Smith T, Willison K, Leaver CJ, Martin W. 2003. The function of genomes in bioenergetic organelles. In: Philosophical Transactions of the Royal Society B: Biological Sciences.

Barreto FS, Watson ET, Lima TG, Willett CS, Edmands S, Li W, Burton RS. 2018. Genomic signatures of mitonuclear coevolution across populations of Tigriopus californicus. Nat Ecol Evol 2:1250–1257.

Barrientos A, Fontanesi F, Díaz F. 2009. Evaluation of the Mitochondrial Respiratory Chain and Oxidative Phosphorylation System Using Polarography and Spectrophotometric Enzyme Assays. Curr Protoc Hum Genet 63.

Barrientos A, Kenyon L, Moraes CT. 1998. Human Xenomitochondrial Cybrids. Journal of Biological Chemistry 273:14210–14217.

Bendich AJ. 1996. Structural Analysis of Mitochondrial DNA Molecules from Fungi and Plants Using Moving Pictures and Pulsed-field Gel Electrophoresis.

Bendich AJ. 2007. The size and form of chromosomes are constant in the nucleus, but highly variable in bacteria, mitochondria and chloroplasts. BioEssays 29:474–483.

Bernardi G. 2005. Lessons from a small, dispensable genome: The mitochondrial genome of yeast. Gene 354:189–200.

Blount ZD, Borland CZ, Lenski RE. 2008. Historical contingency and the evolution of a key innovation in an experimental population of Escherichia coli. Proc Natl Acad Sci U S A 105:7899–7906.

Breslow DK, Cameron DM, Collins SR, Schuldiner M, Stewart-Ornstein J, Newman HW, Braun S, Madhani HD, Krogan NJ, Weissman JS. 2008. A comprehensive strategy enabling high-resolution functional analysis of the yeast genome. Nat Methods 5:711–718.

Breton S, Beaupré HD, Stewart DT, Piontkivska H, Karmakar M, Bogan AE, Blier PU, Hoeh WR. 2009. Comparative mitochondrial genomics of freshwater mussels (Bivalvia: Unionoida) with doubly uniparental inheritance of mtDNA: Gender-specific open reading frames and putative origins of replication. Genetics 183:1575–1589.

Bull JJ, Molineux IJ. 2008. Predicting evolution from genomics: experimental evolution of bacteriophage T7. Heredity (Edinb) [Internet] 100:453–463. Available from: http://www.nature.com/hdy

Burton RS. 2022. The role of mitonuclear incompatibilities in allopatric speciation. Cellular and Molecular Life Sciences 79:103.

Burton RS, Barreto FS. 2012. A disproportionate role for mtDNA in Dobzhansky-Muller incompatibilities? Mol Ecol 21:4942–4957.

Caudal E, Friedrich A, Jallet A, Garin M, Hou J, Schacherer J. 2022. Loss-of-function mutation survey revealed that genes with background-dependent fitness are rare and functionally related in yeast. Available from: https://doi.org/10.1073/pnas.2204206119

Charron G, Marsit S, Hénault M, Martin H, Landry CR. 2019. Spontaneous whole-genome duplication restores fertility in interspecific hybrids. Nat Commun 10.

Chen P, Michel AH, Zhang J. 2022. Transposon insertional mutagenesis of diverse yeast strains suggests coordinated gene essentiality polymorphisms. Nat Commun 13.

Chou HH, Chiu HC, Delaney NF, Segrè D, Marx CJ. 2011. Diminishing returns epistasis among beneficial mutations decelerates adaptation. Science (1979) 332:1190–1192.

Chou JY, Hung YS, Lin KH, Lee HY, Leu JY. 2010. Multiple molecular mechanisms cause reproductive isolation between three yeast species. PLoS Biol.

Cooper CE, Nicholls P, Freedman JA. 1991. Cytochrome c oxidase: structure, function, and membrane topology of the polypeptide subunits. Biochemistry and Cell Biology 69:586–607.

Cooper TF, Rozen DE, Lenski RE. 2003. Parallel changes in gene expression after 20, 000 generations of evolution in Escherichia coli. Proc Natl Acad Sci U S A 100:1072–1077.

Couce A, Tenaillon OA. 2015. The rule of declining adaptability in microbial evolution experiments. Front Genet 6.

Desai N, Brown A, Amunts A, Ramakrishnan V. 2017. The structure of the yeast mitochondrial ribosome. Science (1979) 355:528–531.

Dey R, Barrientos A, Moraes CT. 2000a. Functional constraints of nuclear mitochondrial DNA interactions in xenomitochondrial rodent cell lines. Journal of Biological Chemistry.

Dey R, Barrientos A, Moraes CT. 2000b. Functional Constraints of Nuclear-Mitochondrial DNA Interactions in Xenomitochondrial Rodent Cell Lines. Journal of Biological Chemistry 275:31520–31527.

Dowling DK. 2014. Mitonuclear interactions: evolutionary consequences over multiple biological scales. Philos Trans R Soc Lond B Biol Sci.

Dyall SD, Brown MT, Johnson PJ. 2004. Ancient Invasions: From Endosymbionts to Organelles. Science (1979).

Ferreira GC. 1995. Heme biosynthesis: Biochemistry, molecular biology, and relationship to disease. J Bioenerg Biomembr 27:147–150.

Freel KC, Friedrich A, Schacherer J. 2015. Mitochondrial genome evolution in yeasts: An all-encompassing view. FEMS Yeast Res.

Fritsch E, Chabbert C, Klaus B, Steinmetz L. 2014. A Genome-Wide Map of Mitochondrial DNA Recombination in Yeast. Genetics 198:755–771.

Fritsch ES, Chabbert CD, Klaus B, Steinmetz LM. 2014. A genome-wide map of mitochondrial DNA recombination in yeast. Genetics.

Galeota-Sprung B, Fernandez A, Sniegowski P. 2022. Changes to the mtDNA copy number during yeast culture growth. R Soc Open Sci [Internet] 9. Available from: https://royalsocietypublishing.org/doi/10.1098/rsos.211842

Gueldener U, Heinisch J, Koehler GJ, Voss D, Hegemann JH. 2002. A second set of loxP marker cassettes for Cre-mediated multiple gene knockouts in budding yeast. Nucleic Acids Res [Internet] 30:e23. Available from: http://www.pubmedcentral.nih.gov/articlerender.fcgi?artid=101367&tool=pmcentrez&rendertype=abstract

Haddad R, Meter B, Ross JA. 2018. The Genetic Architecture of Intra-Species Hybrid Mito-Nuclear Epistasis. Front Genet 9:1–11.

Hagström E, Freyer C, Battersby BJ, Stewart JB, Larsson NG. 2014. No recombination of mtDNA after heteroplasmy for 50 generations in the mouse maternal germline. Nucleic Acids Res.

Havird JC, Sloan DB. 2016. The roles of mutation, selection, and expression in determining relative rates of evolution in mitochondrial versus nuclear genomes. Mol Biol Evol.

Hendrich L, Pons J, Ribera I, Balke M. 2010. Mitochondrial Cox1 Sequence Data Reliably Uncover Patterns of Insect Diversity But Suffer from High Lineage-Idiosyncratic Error Rates. PLoS One.

Herron MD, Doebeli M. 2013. Parallel Evolutionary Dynamics of Adaptive Diversification in Escherichia coli. PLoS Biol 11.

Hill GE. 2015. Mitonuclear ecology. Mol Biol Evol.

Hunte C, Palsdottir H, Trumpower BL. 2003. Protonmotive pathways and mechanisms in the cytochrome bc1 complex. FEBS Lett 545:39–46.

Hunter-Manseau F, Desrosiers V, le François NR, Dufresne F, Detrich HW, Nozais C, Blier PU. 2019. From Africa to Antarctica: Exploring the Metabolism of Fish Heart Mitochondria Across a Wide Thermal Range. Front Physiol 10.

Jordá T, Puig S. 2020. Regulation of Ergosterol Biosynthesis in Saccharomyces cerevisiae. Genes (Basel) 11:795.

Kenyon L, Moraes CT. 2002. Expanding the functional human mitochondrial DNA database by the establishment of primate xenomitochondrial cybrids. Proceedings of the National Academy of Sciences.

Kim DU, Hayles J, Kim D, Wood V, Park HO, Won M, Yoo HS, Duhig T, Nam M, Palmer G, et al. 2010. Analysis of a genome-wide set of gene deletions in the fission yeast Schizosaccharomyces pombe. Nat Biotechnol 28:617–623.

Kvitek DJ, Sherlock G. 2013. Whole Genome, Whole Population Sequencing Reveals That Loss of Signaling Networks Is the Major Adaptive Strategy in a Constant Environment. PLoS Genet 9.

Kwan EX, Wang XS, Amemiya HM, Brewer BJ, Raghuraman MK. 2016. rDNA Copy Number Variants Are Frequent Passenger Mutations in Saccharomyces cerevisiae Deletion Collections and de Novo Transformants. G3 Genes|Genomes|Genetics 6:2829–2838.

Lane N. 2011. Mitonuclear match: Optimizing fitness and fertility over generations drives ageing within generations. BioEssays.

Lane N, Martin W. 2010. The energetics of genome complexity. Nature.

Lang GI, Botstein D, Desai MM. 2011. Genetic Variation and the Fate of Beneficial Mutations in Asexual Populations. Genetics 188:647–661.

Lang GI, Rice DP, Hickman MJ, Sodergren E, Weinstock GM, Botstein D, Desai MM. 2013. Pervasive genetic hitchhiking and clonal interference in forty evolving yeast populations. Nature 500:571–574.

Lee HY, Chou JY, Cheong L, Chang NH, Yang SY, Leu JY. 2008. Incompatibility of Nuclear and Mitochondrial Genomes Causes Hybrid Sterility between Two Yeast Species. Cell.

Lenski RE. 2017. Experimental evolution and the dynamics of adaptation and genome evolution in microbial populations. Nature Publishing Group [Internet] 11:2181–2194. Available from: http://www.nature.com/ismej

Linder RA, Greco JP, Seidl F, Matsui T, Ehrenreich IM. 2017. The Stress-Inducible Peroxidase TSA2 Underlies a Conditionally Beneficial Chromosomal Duplication in Saccharomyces cerevisiae. G3 Genes|Genomes|Genetics 7:3177–3184.

Łuksza M, Lässig M. 2014. A predictive fitness model for influenza. Nature 507.

Lynch M, Koskella B, Schaack S. 2006. Mutation pressure and the evolution of organelle genomic architecture. Science (1979).

Lynch M, Sung W, Morris K, Coffey N, Landry CR, Dopman EB, Dickinson WJ, Okamoto K, Kulkarni S, Hartl DL, et al. 2008. A genome-wide view of the spectrum of spontaneous mutations in yeast. Proceedings of the National Academy of Sciences 105:9272–9277.

Ma H, Marti Gutierrez N, Morey R, van Dyken C, Kang E, Hayama T, Lee Y, Li Y, Tippner-Hedges R, Wolf DP, et al. 2016. Incompatibility between Nuclear and Mitochondrial Genomes Contributes to an Interspecies Reproductive Barrier. Cell Metab [Internet] 24:283–294. Available from: http://dx.doi.org/10.1016/j.cmet.2016.06.012

Malina C, Larsson C, Nielsen J. 2018. Yeast mitochondria: an overview of mitochondrial biology and the potential of mitochondrial systems biology. FEMS Yeast Res 18.

Mangin M, Faugeron-Fonty G, Bernardi G. 1983. The orir to ori+ mutation in spontaneous yeast petites is accompanied by a drastic change in mitochondrial genome replication. Gene 24:73–81.

Mason PA, Lightowlers RN. 2003. Why do mammalian mitochondria possess a mismatch repair activity? FEBS Lett.

Mcdonald MJ. 2019. Microbial Experimental Evolution-a proving ground for evolutionary theory and a tool for discovery. EMBO Rep 20.

Mcdonald MJ, Gehrig SM, Meintjes PL, Zhang X-X, Rainey PB. 2009. Adaptive Divergence in Experimental Populations of Pseudomonas fluorescens. IV. Genetic Constraints Guide Evolutionary Trajectories in a Parallel Adaptive Radiation. Genetics [Internet] 183:1041–1053. Available from: http://www.genetics.org/

McKenzie M, Trounce I. 2000. Expression of Rattus norvegicus mtDNA in Mus musculus cells results in multiple respiratory chain defects. Journal of Biological Chemistry.

Meyer JR, Dobias DT, Weitz JS, Barrick JE, Quick RT, Lenski RE. 2012. Repeatability and Contingency in the Evolution of a Key Innovation in Phage Lambda. Science (1979) 335:428–432.

Nguyen THM, Sondhi S, Ziesel A, Paliwal S, Fiumera HL. 2020. Mitochondrial-nuclear coadaptation revealed through mtDNA replacements in Saccharomyces cerevisiae. BMC Evol Biol 20.

Niki Y, Chigusa SI, Matsuura ET. 1989. Complete replacement of mitochondrial DNA in Drosophila. Nature.

Osuský M, Kissová J, Kováč L. 1997. Interspecies transplacement of mitochondria in yeasts. Curr Genet.

Paliwal S, Fiumera AC, Fiumera HL. 2014. Mitochondrial-nuclear epistasis contributes to phenotypic variation and coadaptation in natural isolates of saccharomyces cerevisiae. Genetics.

Puddu F, Herzog M, Selivanova A, Wang S, Zhu J, Klein-Lavi S, Gordon M, Meirman R, Millan-Zambrano G, Ayestaran I, et al. 2019. Genome architecture and stability in the Saccharomyces cerevisiae knockout collection. Nature 573:416–420.

Read AD, Bentley RET, Archer SL, Dunham-Snary KJ. 2021. Mitochondrial iron– sulfur clusters: Structure, function, and an emerging role in vascular biology. Redox Biol 47:102164.

Robba L, Russell SJ, Barker GL, Brodie J. 2006. Assessing the use of the mitochondrial cox1 marker for use in DNA barcoding of red algae (Rhodophyta). Am J Bot.

Rutter J, Hughes AL. 2015. Power2: The power of yeast genetics applied to the powerhouse of the cell. Trends in Endocrinology & Metabolism 26:59–68.

Spinazzi M, Casarin A, Pertegato V, Salviati L, Angelini C. 2012. Assessment of mitochondrial respiratory chain enzymatic activities on tissues and cultured cells. Nat Protoc 7:1235–1246.

Špírek M, Horváth A, Piškur J, Sulo P. 2000. Functional co-operation between the nuclei of Saccharomyces cerevisiae and mitochondria from other yeast species. Curr Genet.

Spirek M, Polakova S, Jatzova K, Sulo P. 2015. Post-zygotic sterility and cytonuclear compatibility limits in S. cerevisiae xenomitochondrial cybrids. Front Genet 5.

Tenaillon O, Rodriguez-Verdugo A, Gaut RL, McDonald P, Bennett AF, Long AD, Gaut BS. 2012. The Molecular Diversity of Adaptive Convergence. Science 335:457–461.

Vijayraghavan S, Kozmin SG, Strope PK, Skelly DA, Lin Z, Kennell J, Magwene PM, Dietrich FS, McCusker JH. 2019. Mitochondrial Genome Variation Affects Multiple Respiration and Nonrespiration Phenotypes in Saccharomyces cerevisiae. Genetics 211:773–786.

Weinreich DM, Delaney NF, DePristo MA, Hartl DL. 2006. Darwinian Evolution Can Follow Only Very Few Mutational Paths to Fitter Proteins. Science (1979) [Internet] 312:111–114. Available from: http://www.sciencemag.org/content/312/5770/111.abstract\n http://www.sciencemag.org/content/312/5770/111.full.pdf

Wichman HA, Badgett MR, Scott LA, Boulianne CM, Bull JJ. 1999. Different Trajectories of Parallel Evolution During Viral Adaptation. Science (1979) 285:422–424.

Wiser M, Ribeck N, Lenski R. 2013. Long-term dynamics of adaptation in asexual populations. Science (1979) 342:1364–1367.

Wolters JF, Charron G, Gaspary A, Landry CR, Fiumera AC, Fiumera HL. 2018. Mitochondrial recombination reveals mito–mito epistasis in yeast. Genetics 209:307–319.

Wolters JF, Chiu K, Fiumera HL. 2015. Population structure of mitochondrial genomes in Saccharomyces cerevisiae. BMC Genomics [Internet] 16:1–13. Available from: http://dx.doi.org/10.1186/s12864-015-1664-4

Xu F, Addis JBL, Cameron JM, Robinson BH. 2012. LRPPRC mutation suppresses cytochrome oxidase activity by altering mitochondrial RNA transcript stability in a mouse model. Biochemical Journal 441:275–283.

Yamaoka M, Isobe K, Shitara H, Yonekawa H, Miyabayashi S, Hayashi JI. 2000. Complete repopulation of mouse mitochondrial DNA-less cells with rat mitochondrial DNA restores mitochondrial translation but not mitochondrial respiratory function. Genetics.

Zakharov I, Yarovoy B. 1977. Cytoduction as a new tool in studying cytoplasmic heredity in yeast. Mol Cell Biochem 14:15–18.

de Zamaroczy M, Marotta R, Faugeron-Fonty G, Coursot R, Mangin M, Baldacci G, Bernardi G. 1981. The origins of replication of the yeast mitochondrial genome and the phenomenon of suppressivity. Nature 292:75–78.

