## Supplementary figures for "Evolutionary trajectories are contingent on mitonuclear interactions"

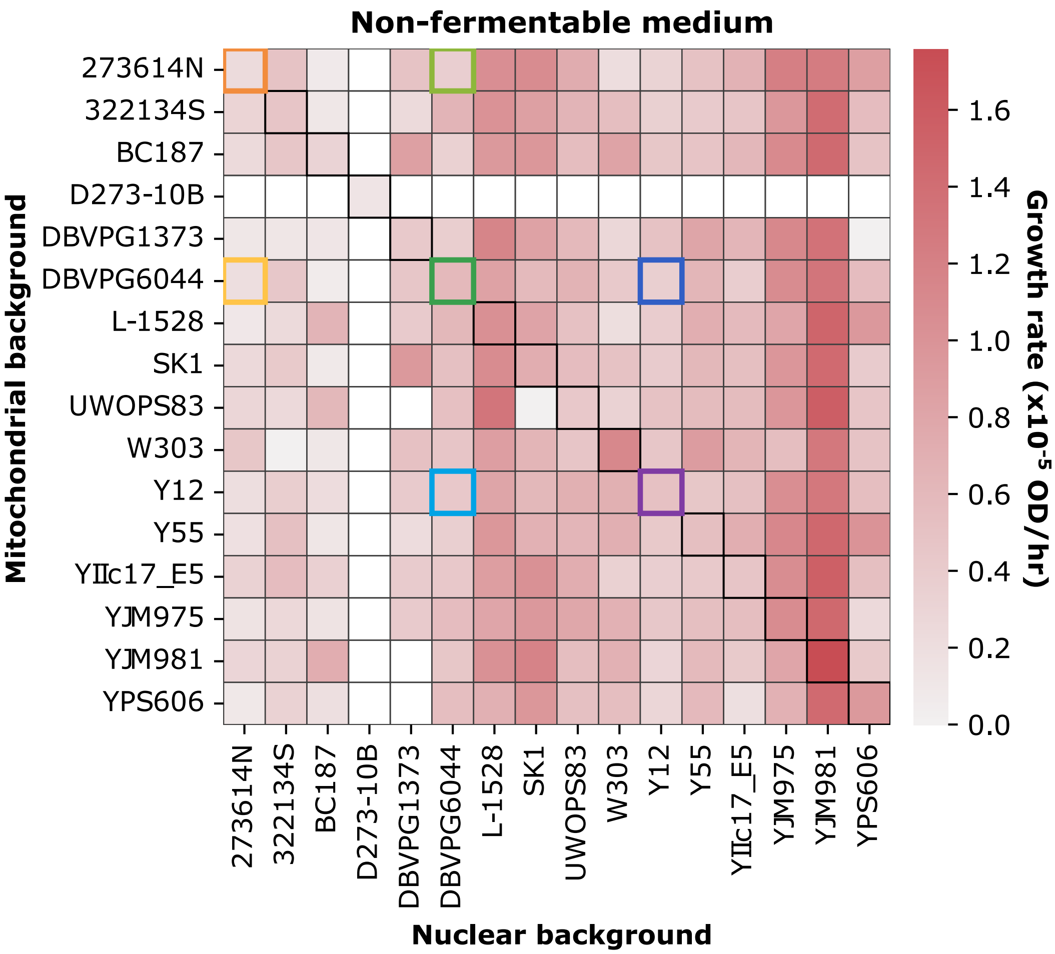


**Figure S1. Screening of a collection of yeast cybrids identifies examples of mitonuclear negative epistasis on fitness**. Growth rate was measured in non-fermentable medium for all individuals in a collection of 225 mitonuclear backgrounds, representing all combinations of nuclear and mitochondrial genomes from a set of 16 yeast strains. Among those, cybrids derived from crosses of strain DBVPG6044 with strains 273614N and Y12 (indicated by colored boxes) appeared to display a reduction in fitness, hinting at mitonuclear incompatibilities. Those strains were submitted to further characterization, and later experimental evolution.

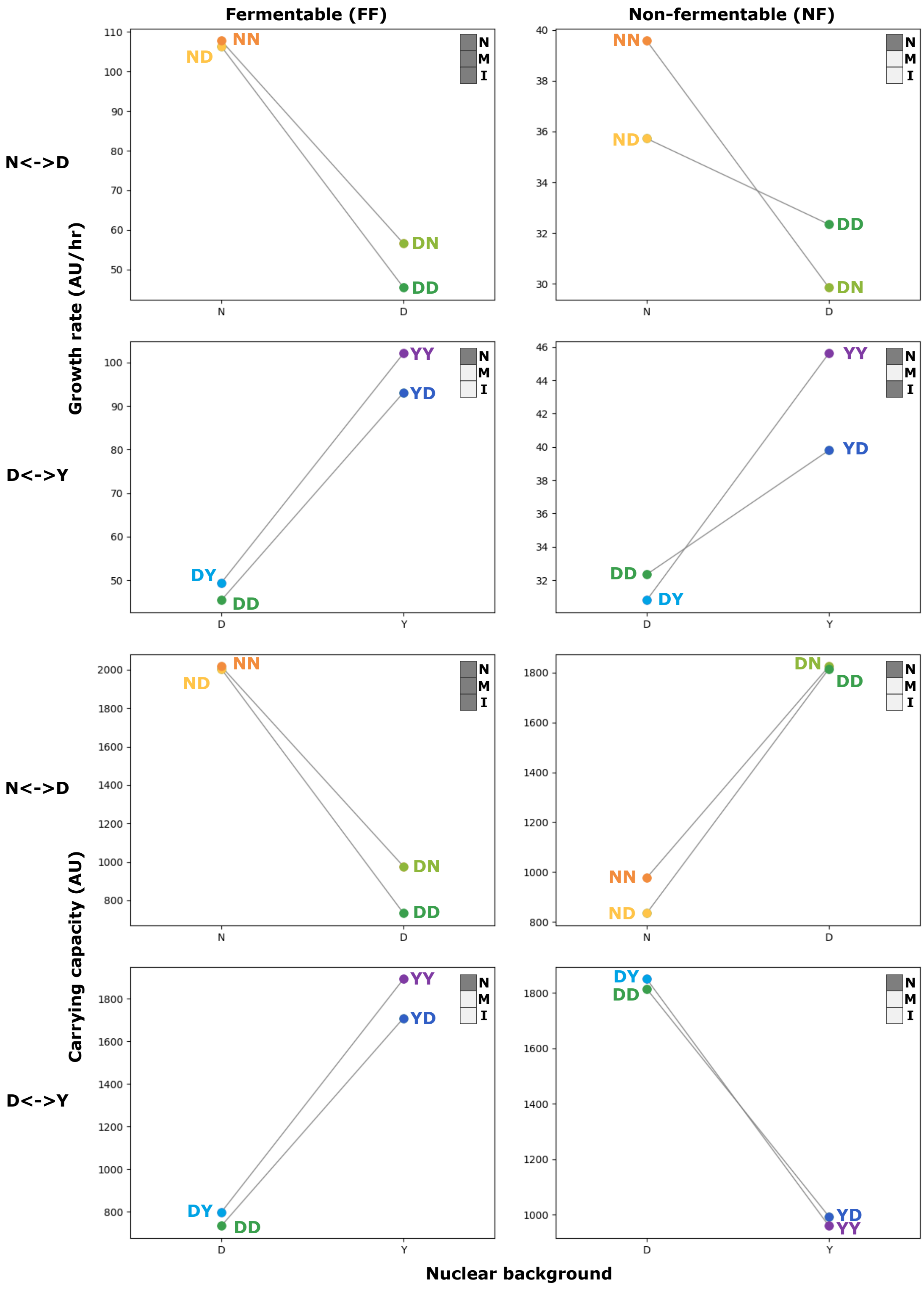
Fermentable (FF)
Non-fermentable (NF)
N<->D
D<->Y
N<->D
D<->Y
Growth rate (AU/hr)
Carrying capacity (AU)
Nuclear background
NN
ND
DN
DD
DY
YY
YD
N
M
I

**Figure S2. Growth parameters of founding strains of experimental evolution reveal nuclear, mitochondrial and mitonuclear effects.** Growth rate (top four panels) and carrying capacity (bottom four panels) measured for all seven strains in fermentable (left) and non-fermentable (right) media are reported as interaction plots for crosses of NN with DD (N<->D) and DD with YY (D<->Y). In each panel, upper right corner insets indicate if nuclear (N) and mitochondrial (M) backgrounds, as well as mitonuclear interactions (I) have a significant effect on growth rate (two-way ANOVA p-value < 0.05). Plots report the mean values of four biological replicates, estimated from 8 to 42 technical replicates, for an average of 27 per point.

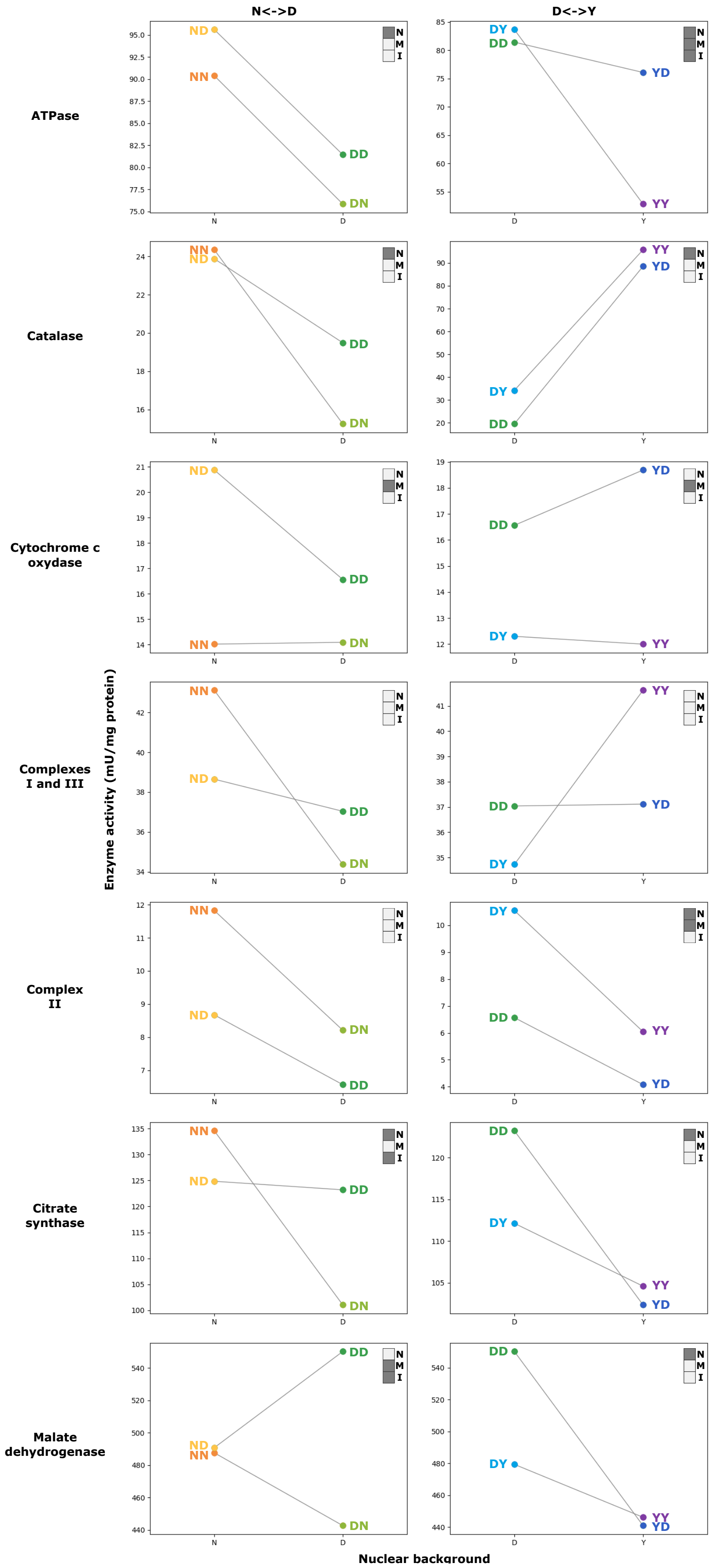
N<->D
D<->Y
ATPase
Catalase
Cytochrome c oxydase
Complexes I and III
Complex II
Citrate synthase
Malate dehydrogenase
Enzyme activity (mU/mg protein)
Nuclear background
N
M
I
NN
ND
DD
DN
DY
YD
YY

**Figure S3. Activity of enzymes associated with the mitochondrion reveal nuclear, mitochondrial and mitonuclear effects in founding strains of evolution experiment.** Activity of the enzymes indicated on the left-hand side and assayed in whole-cell extracts is reported as interaction plots for crosses of NN with DD (N<->D) and DD with YY (D<->Y). In each panel, upper right corner insets indicate if nuclear (N) and mitochondrial (M) backgrounds, as well as mitonuclear interactions (I) have a significant effect on growth rate (two-way ANOVA p-value < 0.05). Plots report the mean values of three or four biological replicates.

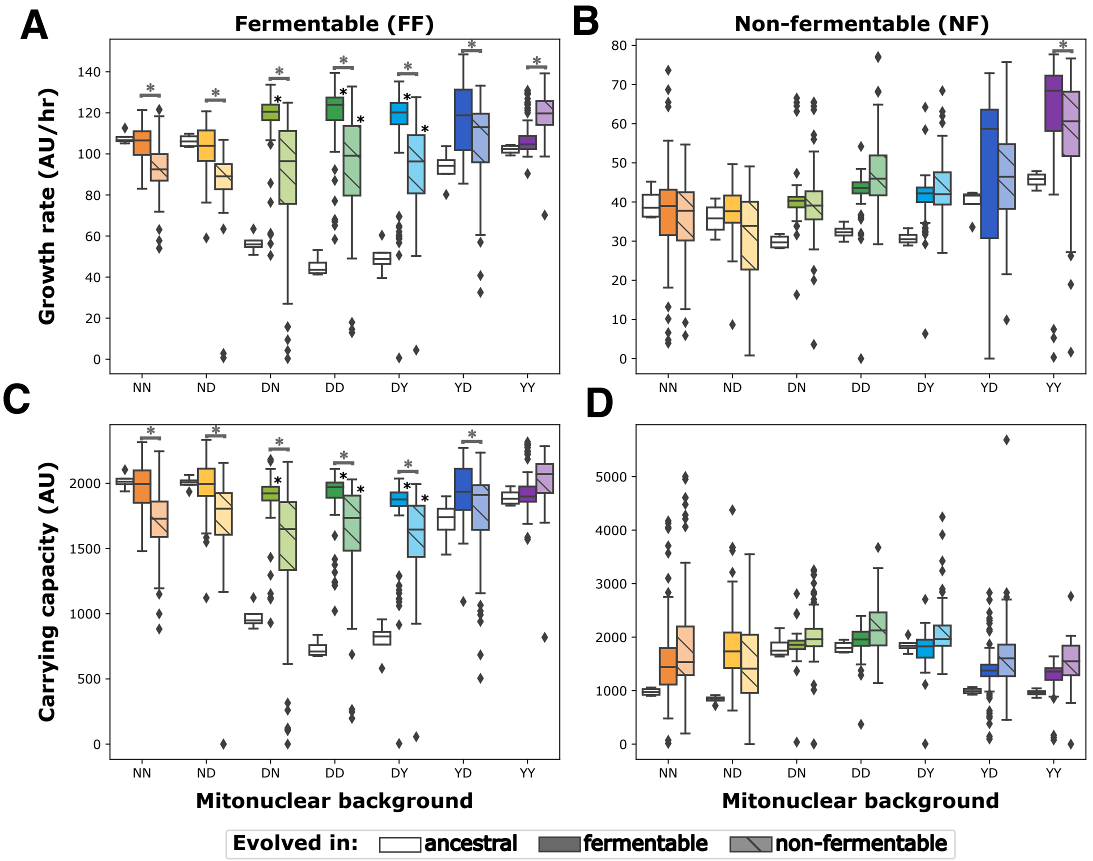


**Figure S4. Experimental evolution of mitonuclear hybrids leads to fitness change in both fermentable and non-fermentable media in a mitonuclear background and environment dependent manner.** Fitness of evolved individuals and their ancestral strains was estimated by measuring growth rate **(A,B)** and carrying capacity **(C,D)** in fermentable **(A,C)** and non-fermentable **(B,D)** media. Two-way ANOVAs indicate significant effects on growth rate for mitonuclear background and evolution regimen. Boxes marked by asterisks (*) indicate evolved strains with mean fitness significantly above ancestral levels (Tukey HSD test p-value < 0.05). Asterisk (*) marked connectors indicate mean fitness significantly different between strains of the same mitonuclear background evolved in different environments (Tukey HSD test p-value < 0.05).

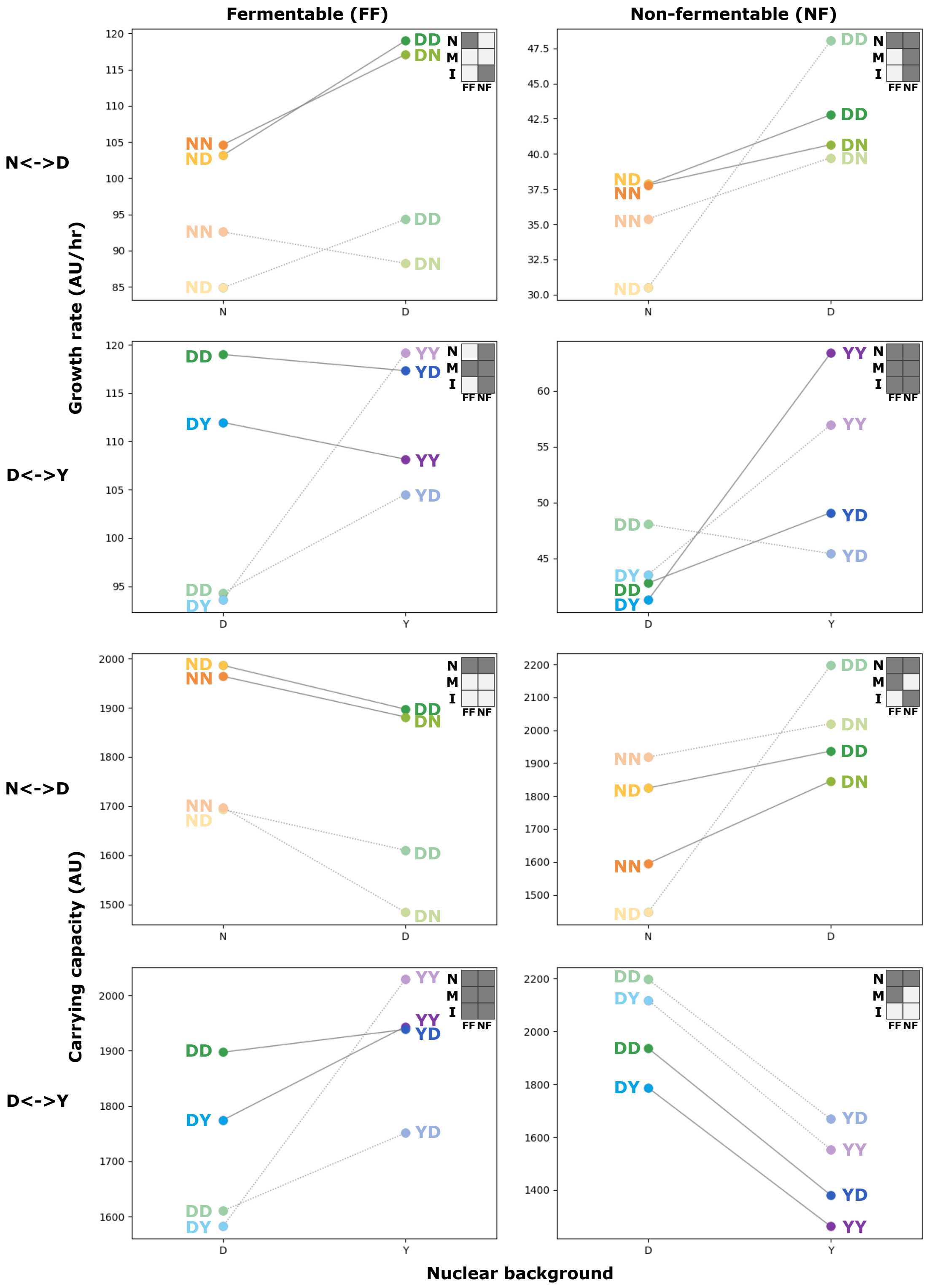
Fermentable (FF)
Non-fermentable (NF)
N<->D
D<->Y
N<->D
D<->Y
Growth rate (AU/hr)
Carrying capacity (AU)
Nuclear background
N
M
I
FF
NF
NN
ND
DD
DN
DY
YY
YD

**Figure S5. Growth parameters of evolved strains reveal nuclear, mitochondrial and mitonuclear effects.** Interaction plots and two-way ANOVAs (upper right insets) indicate significant effects (p-value < 0.05) for nuclear background (N), mitochondrial background (M) or mitonuclear interactions (I) on growth rate (top four panels) or carrying capacity (bottom four panels) in fermentable (left) and non-fermentable (right) media for strains of both crosses evolved in fermentable (dark hues, FF in inset) and non-fermentable (light hues, NF in inset) environments. Plots report the mean value for 37 to 91 strains, for an average of 80 per point.

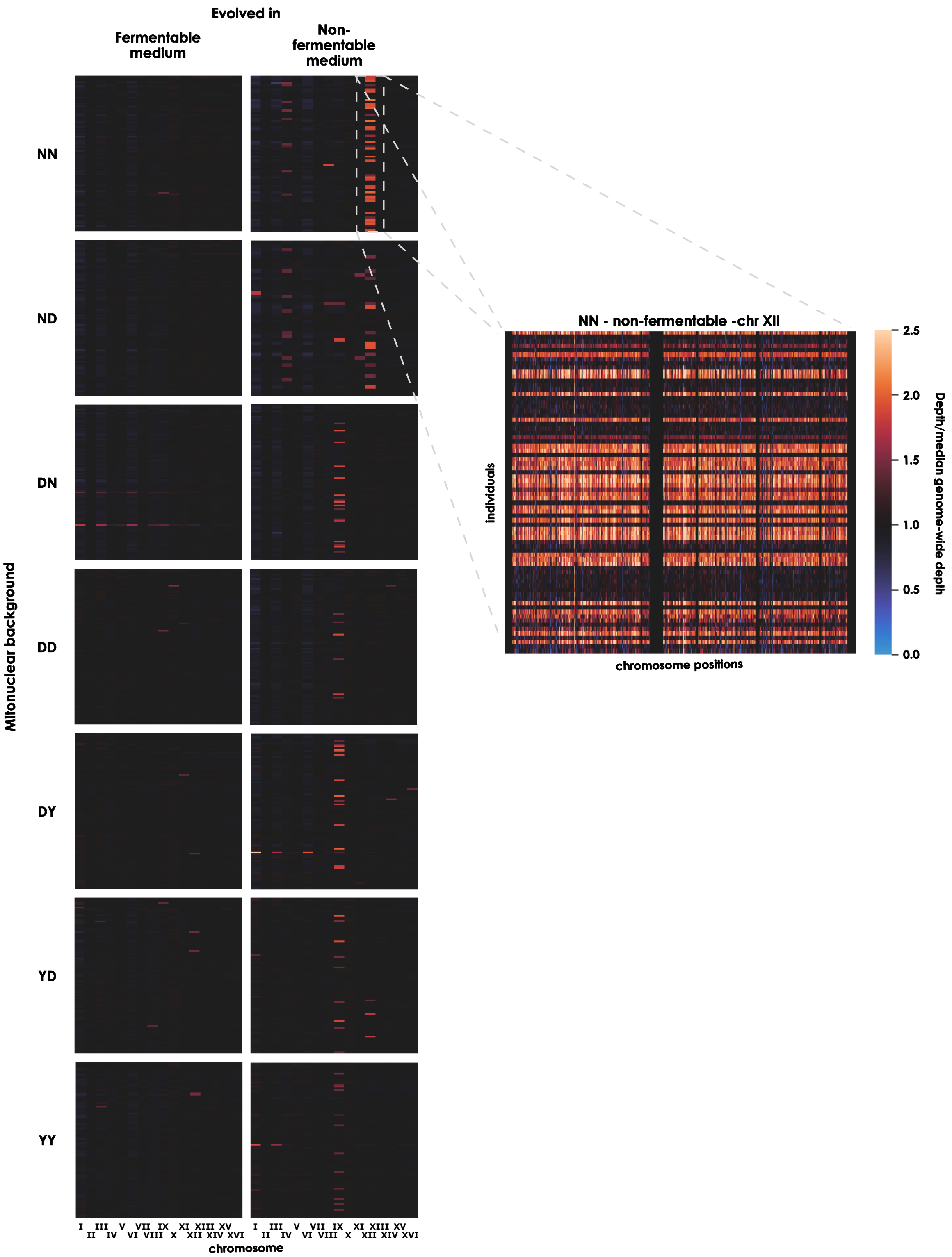
Evolved in
Fermentable medium
Non-fermentable medium
Mitonuclear background
NN
ND
DN
DD
DY
YD
YY
I III V VII IX XI XIII XV
II IV VI VIII X XII XIV XVI
chromosome
NN - non-fermentable -chr XII
Individuals
chromosome positions
Depth/median genome-wide depth
2.5
2.0
1.5
1.0
0.5
0.0

**Figure S6. Visual examination of aneuploidy in evolved individuals.** The ratio of median chromosome depth to chromosome-wide median depth is reported for individuals of each mitonuclear background, evolved in fermentable and non-fermentable media. As an example, detailed depth along chromosome XII is reported for NN individuals evolved in non-fermentable conditions.

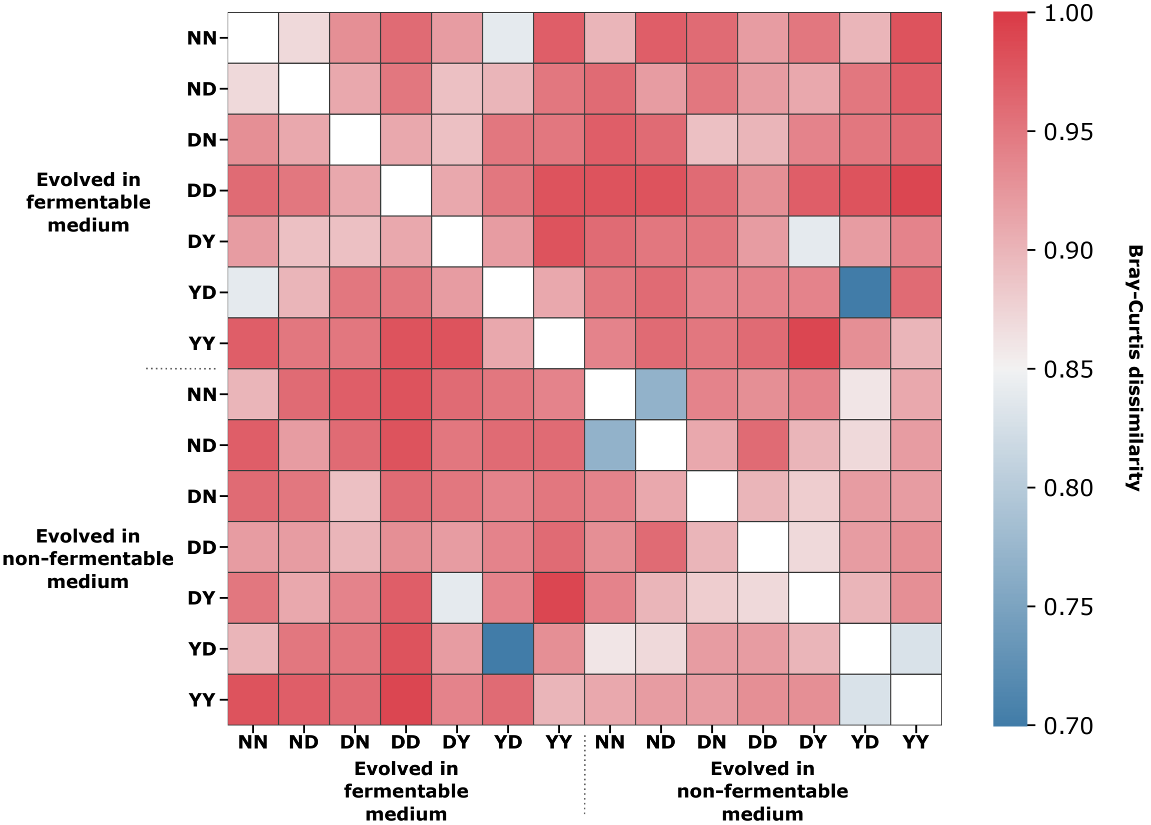


**Figure S7. Dissimilarity between mutational profiles of all mitonuclear backgrounds evolved in both carbon sources.** The heatmap reports Bray-Curtis dissimilarity computed for each pair of evolutionary circumstances as shades of blue (lower dissimilarity) to red (higher dissimilarity). Comparisons of self against self (diagonal) are left blank.

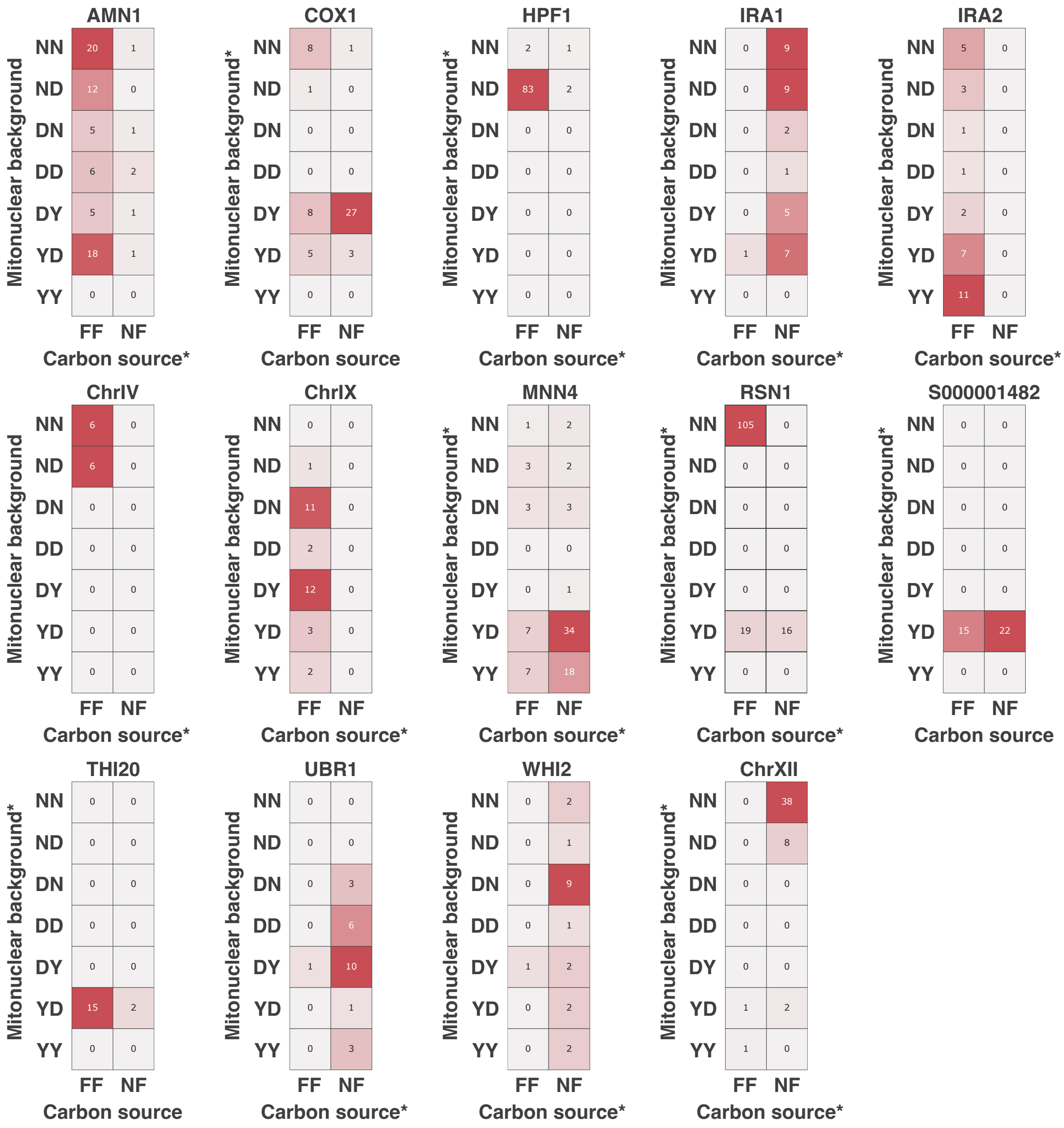


**Figure S8. Patterns of convergent evolution for mutation at certain loci suggests carbon source and mitonuclear specificity.** Partitioning of mutations between mitonuclear backgrounds and carbon sources is shown for the mutations mapping to the indicated annotations.

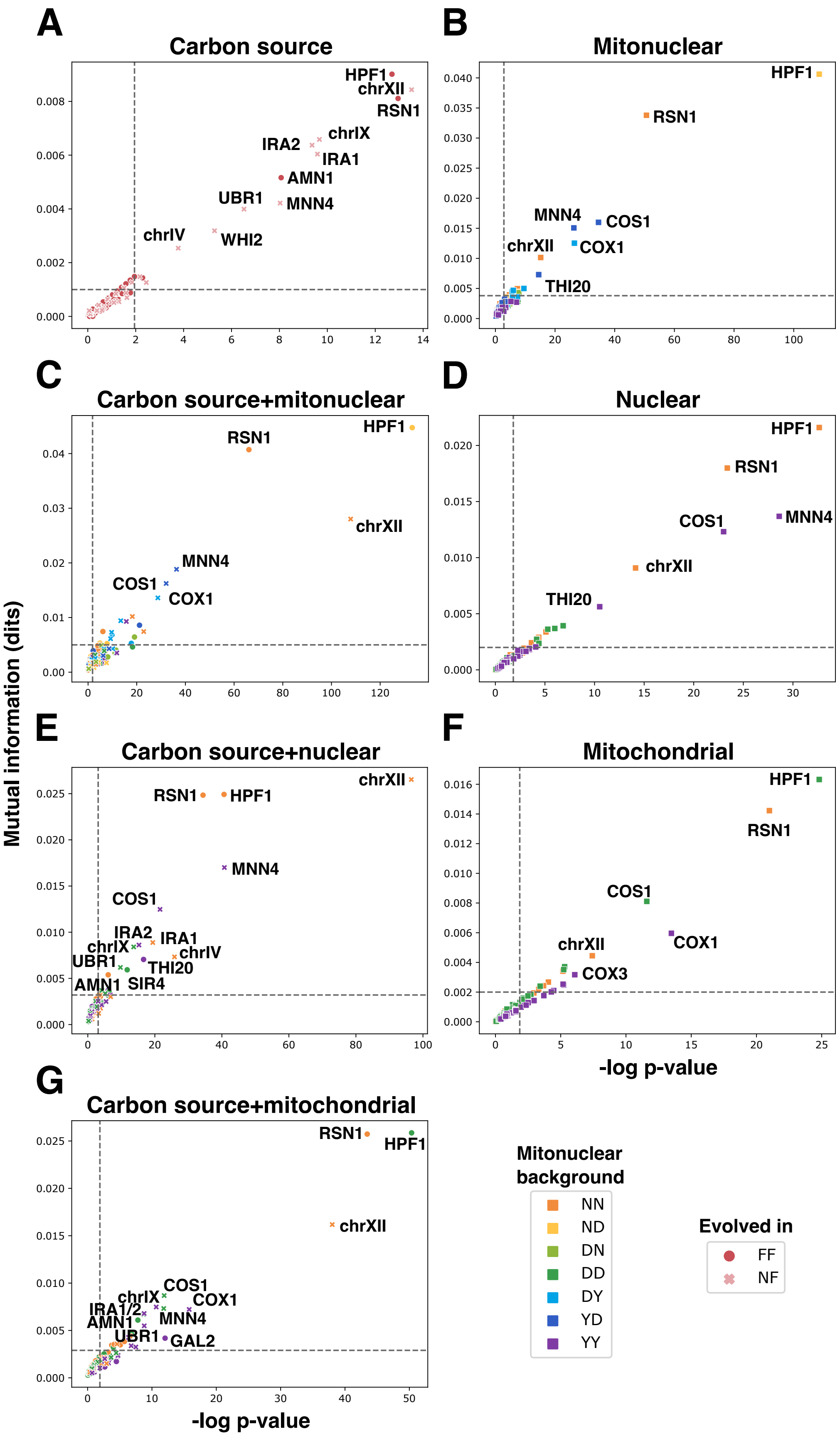
A
Carbon source
B
Mitonuclear
C
Carbon source+mitonuclear
D
Nuclear
E
Carbon source+nuclear
F
Mitochondrial
G
Carbon source+mitochondrial
Mutual information (dits)
-log p-value
HPF1
chrXII
RSN1
chrIX
IRA2
IRA1
AMN1
UBR1
MNN4
chrIV
WHI2
COS1
COX1
THI20
IRA1/2
SIR4
COX3
GAL2
Mitonuclear background
NN
ND
DN
DD
DY
YD
YY
Evolved in
FF
NF

**Figure S9. Mutations at certain loci display specificity for carbon source, nuclear background, mitochondrial background, and various combinations of all three factors.** Specificity for mutations at all loci was tested for carbon source **(A)**, mitonuclear background **(B)**, combination of carbon source and mitonuclear background **(C)**, nuclear background **(D)**, combination of carbon source and nuclear *background* **(E**), mitochondrial background **(F)**, or combination of carbon source and mitochondrial background. For each locus, $chi^2$ tests were performed on contingency tables of all mutations, classifying them as mapping or not to the locus, and according to either carbon source or mitonuclear background. Mutual information was calculated from the same contingency tables. Specificity thresholds, indicated by dotted lines, were set at a p-value of 0.05, corrected for false-discovery rate, and at two standard deviations above mean (~$95^{th}$ percentile) mutual information across all loci.

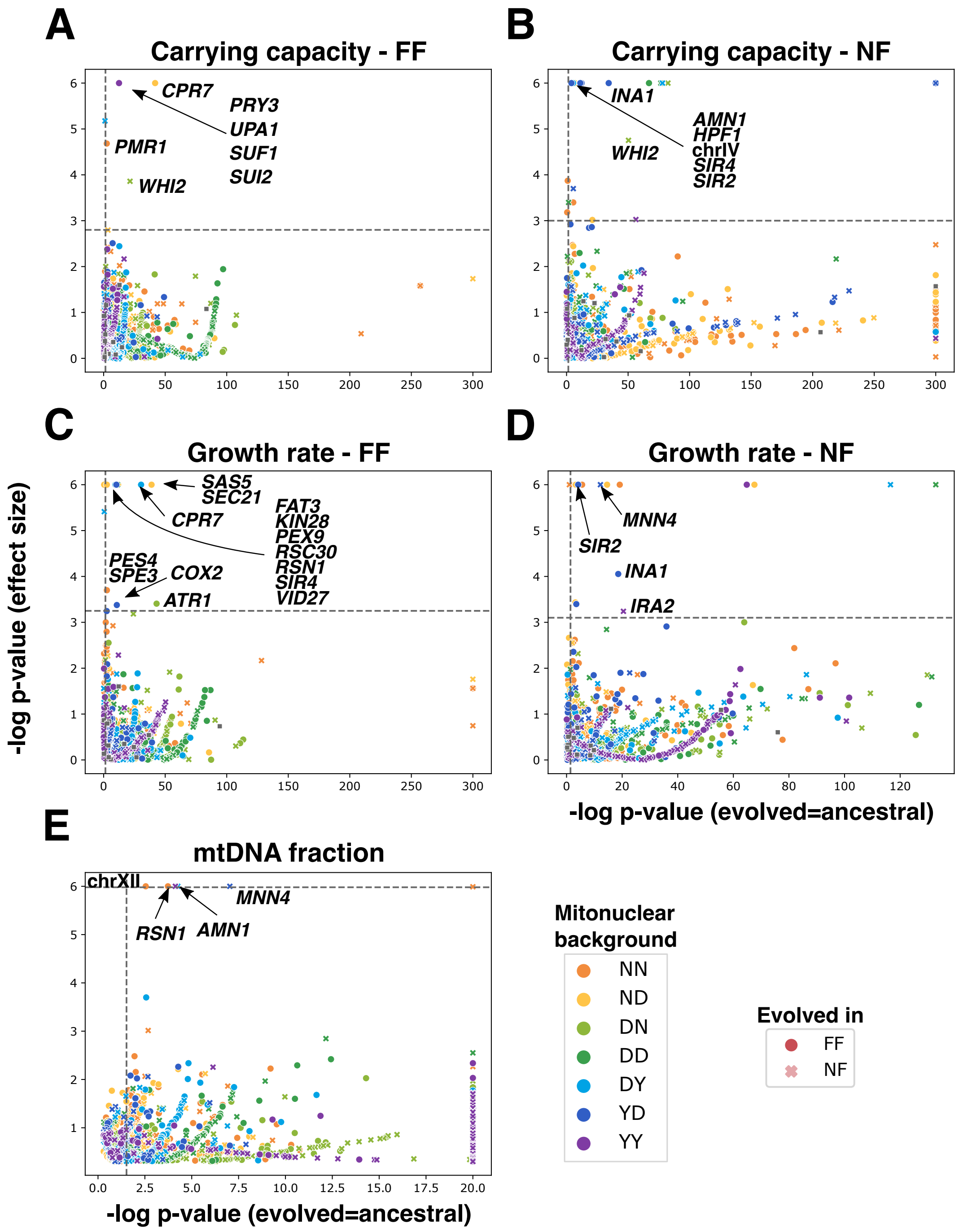
A
Carrying capacity - FF
CPR7
PRY3
UPA1
SUF1
SUI2
PMR1
WHI2
B
Carrying capacity - NF
INA1
AMN1
HPF1
chrIV
SIR4
SIR2
WHI2
C
Growth rate - FF
SAS5
SEC21
CPR7
FAT3
KIN28
PEX9
RSC30
RSN1
SIR4
VID27
PES4
SPE3
COX2
ATR1
D
Growth rate - NF
MNN4
SIR2
INA1
IRA2
E
mtDNA fraction
chrXII
MNN4
RSN1
AMN1
-log p-value (effect size)
-log p-value (evolved=ancestral)
Mitonuclear background
NN
ND
DN
DD
DY
YD
YY
Evolved in
FF
NF

**Figure S10. Mutation at certain loci is associated with quantitative phenotype changes.** Mutant loci associated with quantitative changes in phenotype were identified by performing two statistical tests. Scatter plots show the negative log p-value that individuals mutant at a given locus have phenotype identical to their parents on the x-axis, and the negative p-value that a phenotypic effect of this size could be observed from a set of random individuals on the y-axis. Dotted lines indicate significance thresholds for p-value=0.05, corrected for false-discovery rate. Color of the data points indicates the mitonuclear background most often mutant at the locus, while shape indicates carbon source in which the locus was most often mutant. This analysis was performed for carrying capacity **(A, B)** and growth rate **(C, D)**, in fermentable **(A, C)** and non-*fermentable* **(B, D)** media, as well as for the fraction of mtDNA per cell **(E)**.

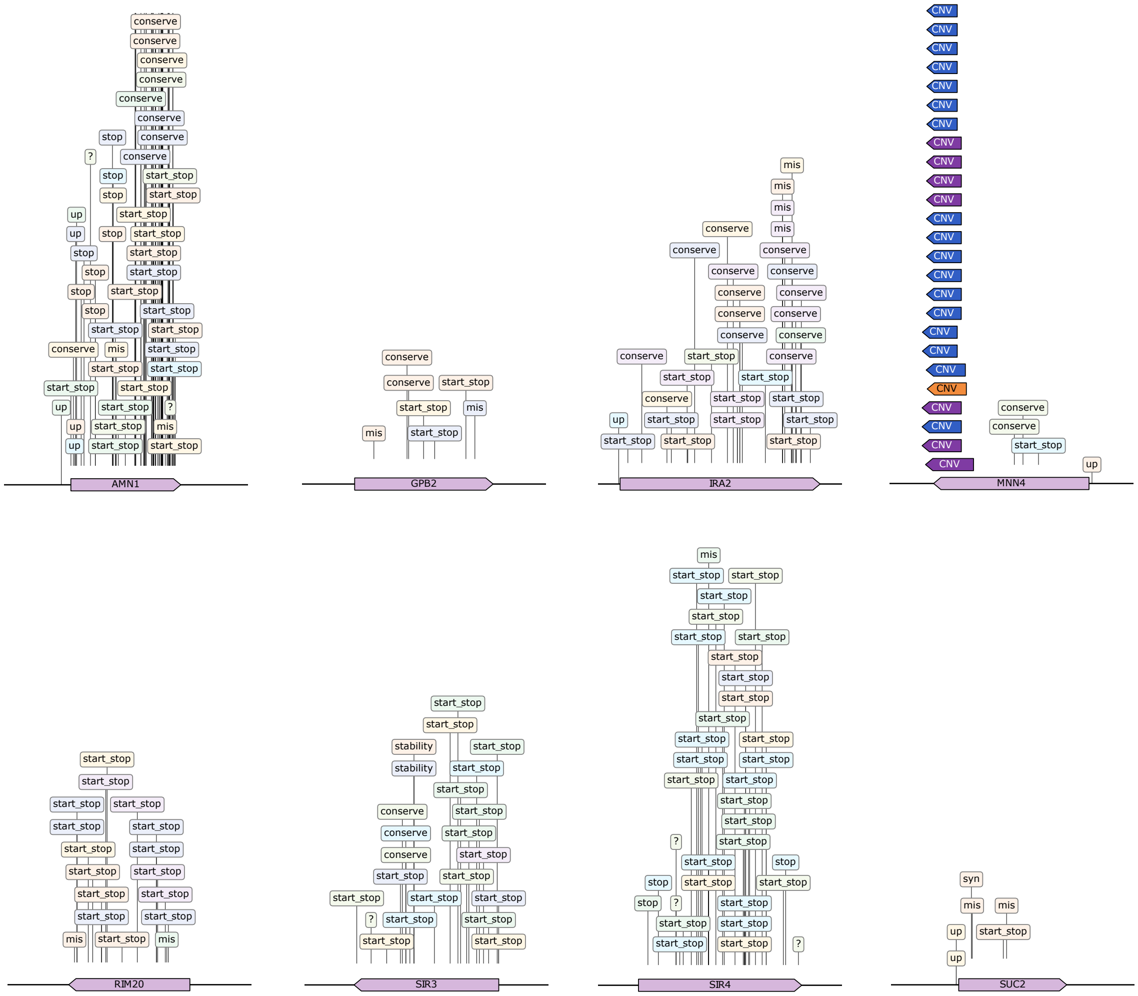


**Figure S11. Predicted effect of mutations detected by sequencing frequently suggests a loss-of-function mechanism.** Effect of mutations was predicted with SNPeff and mapped on the sequence of their locus.”Start-stop” and “stop” signposts indicate nonsense mutations. “mis” indicates missense mutations while “conserved” indicates missense mutations at conserved positions. “up” indicates a mutation upstream of the open reading frame. “CNV” indicates a copy number variant. Question marks (?) are indicative of mutations for which no effect was inferred.

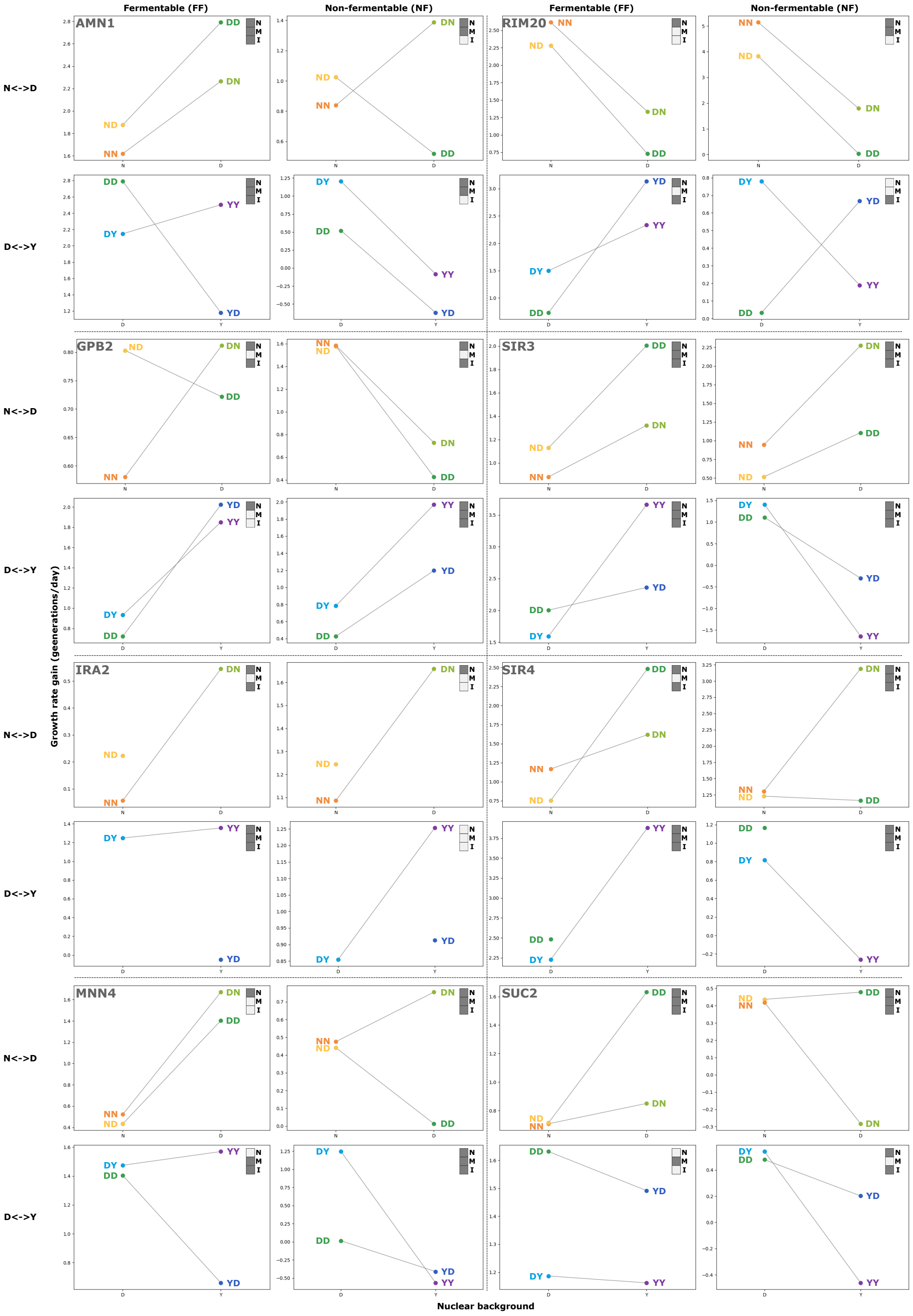
Fermentable (FF)
Non-fermentable (NF)
Fermentable (FF)
Non-fermentable (NF)
AMN1
RIM20
GPB2
SIR3
IRA2
SIR4
MNN4
SUC2
N<->D
D<->Y
N<->D
D<->Y
N<->D
D<->Y
N<->D
D<->Y
Growth rate gain (geenerations/day)
Nuclear background

**Figure S12. Mitonuclear interactions influence fitness gains associated with loss-of-function at loci selected by evolution.** Growth rate gains recorded for all seven strains in fermentable (left) and non-fermentable (right) media are reported as interaction plots for crosses of NN with DD (N<->D) and DD with YY (D<->Y). In each panel, upper right corner insets indicate if nuclear (N) and mitochondrial (M) backgrounds, as well as mitonuclear interactions (I) have a significant effect on gains (two-way ANOVA p-value < 0.05). Dashed lines separate analyses for each of the eight tested loci. Identity of the deleted loci is indicated in the in the upper left corner of each block of interaction plots.

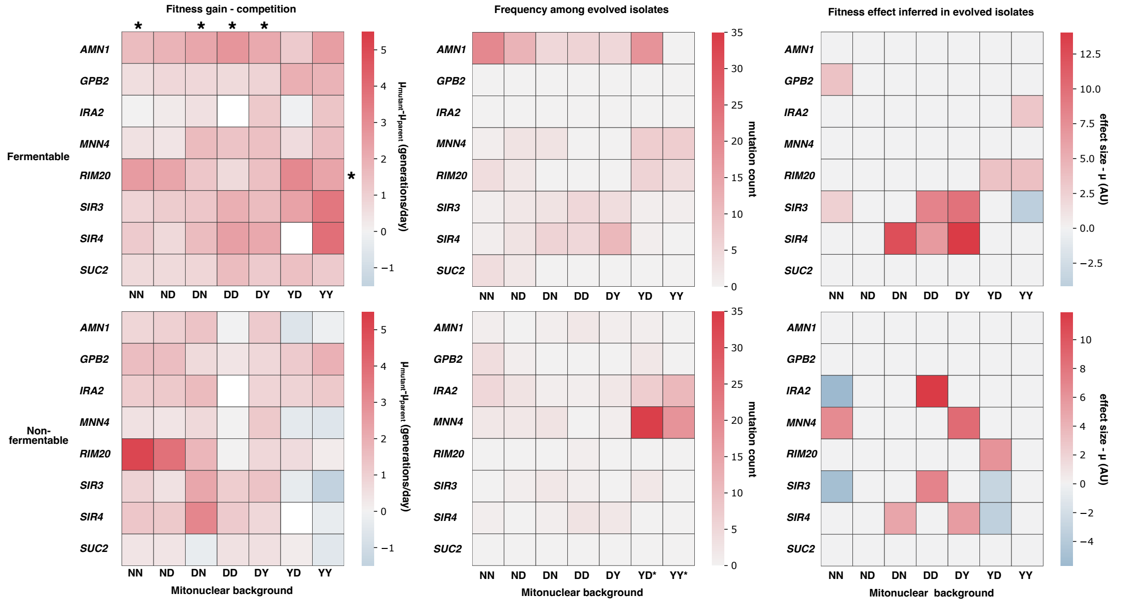


**Figure S13. Fitness effect for loss-of-function at select loci is dependent on mitonuclear background and carbon source.** Gene knockouts were performed at the indicated loci in fluorescent-derivatives of all seven ancestral strains used in the evolution experiment, mimicking the effect of loss-of-function alleles. These knockouts were competed against their deletion-free counterparts of the same mitonuclear genotype **(left panels)** in fermentable **(top)** and non-fermentable **(bottom)** media. This enabled estimation of the fitness effect of loss-of-function at these loci, reported in shades of red (positive effect) or blue (negative effect). For each locus, effect of deletion was found to differ significantly between mitonuclear genotypes (ANOVA p-value < 0.01), as indicated by asterisks (*) next to locus names. Similarly, loci had differing effects on any given mitonuclear genotype (ANOVA p-value < 0.01), as indicated by asterisks (*) next to genotype identifiers. Fitness effect of each locus over mitonuclear genotypes did not correlate with frequency of mutation at the locus in each genotype **(middle panels)**, with one exception indicated by an asterisk on the right-hand side (spearman rho > 0.95). Asterisks at the top of heatmaps indicate genotypes that display rank order correlation between locus fitness effect preference and frequency of mutation at the locus (spearman rho > 0.7, p-value<0.05 that slope of regression =0). Fitness effects estimated from competition assays may be compared to fitness effect inferred from evolved strains carrying the mutations **(right panels)**.

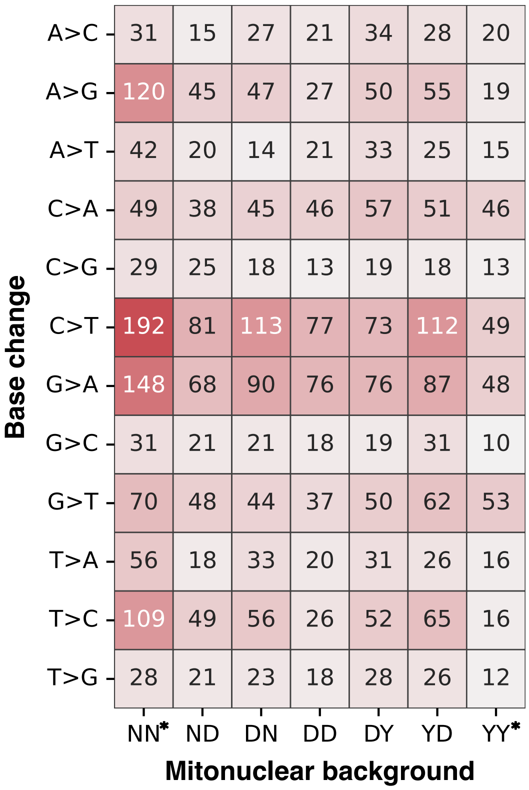


**Figure S14. Single-nucleotide changes detected in evolved individuals are indicative of mutational bias.** Counts of single-nucleotide changes for every possible base change are found for each mitonuclear background. Asterisks indicate backgrounds for which the pattern of mutational bias differs significantly from the rest of the dataset ($chi^2$ test of independence p-value > 0.05).

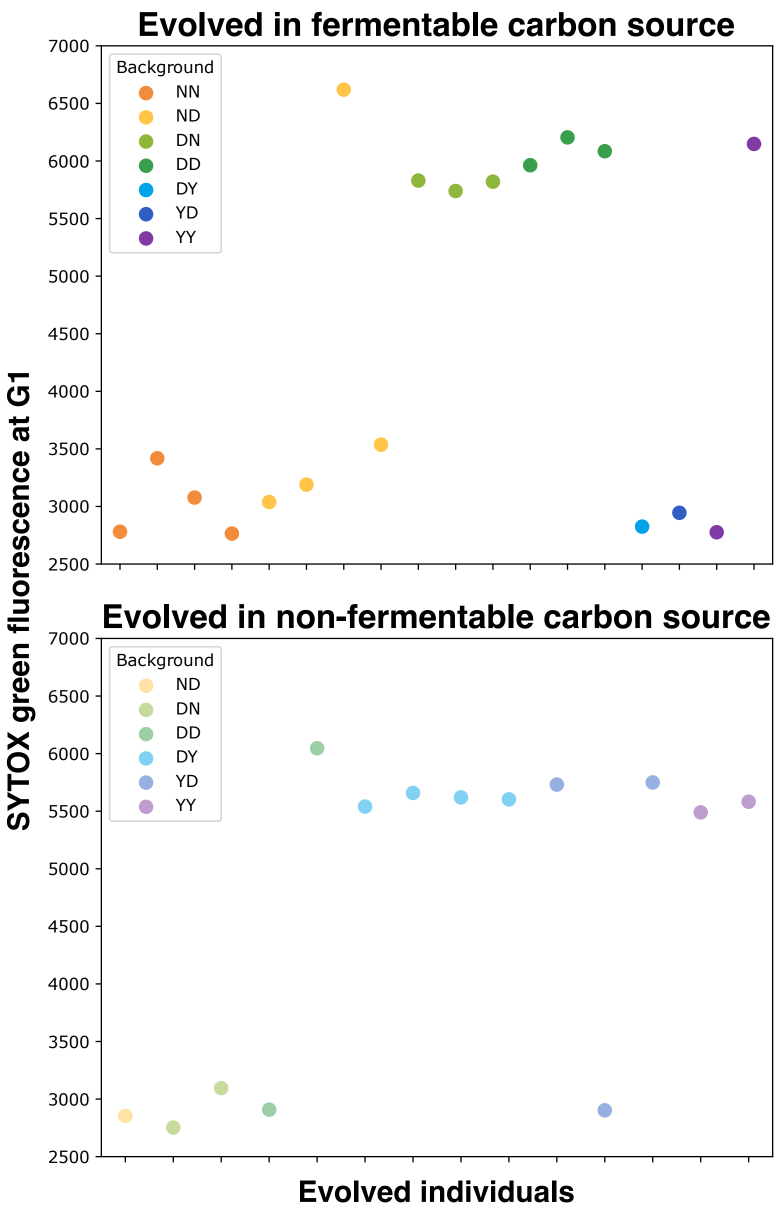


**Figure S15. Experimental evolution often leads to diploidization in ancestrally haploid populations.** Ploidy was assessed in select individuals by SYTOX green staining and flow cytometry. Cells were fixed in ethanol and treated with RNase A before staining nucleic acids with SYTOX green. Stained cells were analyzed by cytometry, visually identifying individuals in G1 phase and measuring their mean SYTOX green-associated fluorescence. High fluorescence is indicative of a change in ploidy.
